# A single mycobacterial ligand organizes multi-receptor signaling to reprogram macrophage lipid metabolism

**DOI:** 10.64898/2026.02.18.706227

**Authors:** Dhrubajyoti Nag, Joycelyn Radeny, Jinyan Cui, Omair Vehra, Yan Yu, Jérôme Nigou, Samantha L. Bell, Maria L Gennaro

## Abstract

How innate immune receptors integrate signals from complex microbial ligands remains poorly understood, yet this integration may offer new avenues for host-directed therapies. Here, we show that the architecture of a single pathogen-derived component can organize the coordinated engagement of multiple pattern-recognition receptors to reprogram host cell behavior. We find that the mycobacterial lipoglycan mannose-capped lipoarabinomannan (ManLAM) uses distinct structural features to engage two pattern-recognition receptors, Toll-like receptor 2 (TLR2) and Dectin-2, thereby driving macrophage lipid remodeling and lipid droplet accumulation, a process linked to foam cell formation and necrotizing lesion development in tuberculosis. Dual receptor engagement also potentiates NF-κB-dependent inflammatory signaling, while lipid droplet accumulation proceeds through an mTORC1–PPARγ-dependent pathway that is largely independent of NF-κB activation, indicating that metabolic and inflammatory programs are mechanistically separable. ManLAM-induced lipid remodeling closely mirrors that induced by *Mycobacterium tuberculosis* infection in both neutral lipid composition and pathway dependence. In contrast, other mycobacterial ligands that are lipogenic in vitro do not measurably contribute to lipid droplet accumulation during infection. These findings identify ManLAM as a major mycobacterial driver of lipid remodeling associated with foam cell formation and establish ligand architecture as a mechanism by which complex microbial ligands organize multi-receptor signaling to direct distinct host cell programs.

**Significance statement:** Our findings establish the principle that the architecture of a single microbial ligand can organize the co-engagement of multiple innate immune receptors to shape host cell responses. Using the mycobacterial lipoglycan mannose-capped lipoarabinomannan as a model, we show that distinct structural features within a single microbial component coordinate the co-engagement of TLR2 and Dectin-2 to reprogram macrophage lipid metabolism and promote lipid droplet formation, a process linked to foam cell formation and necrotizing tuberculosis lesions. These results identify a mechanism by which complex microbial ligands can integrate host sensing pathways through their molecular structure. By defining the receptor-signaling axes that control foam cell formation, this work highlights host pathways as candidate targets for host-directed intervention.

## Introduction

Tuberculosis (TB) remains the leading cause of death worldwide from a single infectious agent (1). Infection with *Mycobacterium tuberculosis* drives the formation of lung granulomas, which, under sustained immune stimulation, can progress from protective structures to necrotizing lesions that underlie lung tissue destruction and transmission (2–4). A defining feature of this transition is the emergence of lipid-droplet-laden macrophage foam cells, which exhibit impaired antimicrobial activity, amplify pro-inflammatory and pro-necrotic signaling, and directly contribute to the necrotic core (4, 5). Foam cells may also provide a lipid-rich environment supporting *M. tuberculosis* persistence and sequester lipophilic antibiotics such as bedaquiline (5, 6). Despite their central role at the interface of immunity, metabolism, and pathology, the mechanisms driving foam cell biogenesis in TB remain poorly defined.

Several mycobacterial components, most of them associated with the cell envelope, have been linked to macrophage lipid droplet accumulation and engage multiple pattern-recognition receptors. How signals from these receptors are coordinated to drive macrophage lipid remodeling remains unclear. Mannose-capped lipoarabinomannan (ManLAM) is the best characterized of these factors. ManLAM is a ∼17-kDa structurally heterogeneous lipoglycan composed of a mannosyl-phosphatidylinositol anchor, a mannan core, and an arabinan domain that is capped with terminal mannose residues in pathogenic mycobacteria, including the *M. tuberculosis* complex (7–9). In addition to its structural role in cell wall integrity (10), ManLAM is a major determinant of mycobacterial virulence, disrupting macrophage phagosome maturation and modulating adaptive immune responses, including CD1-restricted T cell activation, CD4_⁺_ T cell function, and cytokine and antibody production by B cells (8, 9). Such pleiotropic effects reflect ManLAM’s capacity to engage multiple pattern-recognition receptors on macrophages, including C-type lectin receptors (CLRs) such as the mannose receptor and Dectin-2, as well as Toll-like receptor (TLR) 2. Notably, ManLAM binds Dectin-2 with high avidity (11, 12), whereas its interaction with TLR2 occurs with comparatively low affinity (13, 14). Seminal work by D’Avila et al. (15) demonstrated that TLR2 is required for ManLAM-induced lipid droplet accumulation, yet activation of TLR2 alone by a potent agonist fails to induce this response (16). These observations indicate that TLR2 signaling is necessary but not sufficient to promote foam cell formation, leaving unresolved the receptor mechanism by which ManLAM drives lipid droplet biogenesis.

We identify coordinated engagement of TLR2 and Dectin-2 as the mechanistic principle underlying ManLAM-driven foam cell formation. Distinct structural moieties within ManLAM engage these pattern-recognition receptors to activate an mTORC1-PPARγ axis that drives triglyceride synthesis and lipid droplet biogenesis in macrophages. These same structural determinants are also required for full inflammatory activation. Together, these findings establish a mechanistic framework for foam cell formation in TB and identify host pathways that represent potential points of vulnerability for host-directed therapeutic intervention.

## Results

### *M. tuberculosis* ManLAM induces macrophage lipid droplet accumulation through TLR2 and Dectin-2

Previous studies showed that the vaccine strain *Mycobacterium bovis* Bacillus Calmette-Guérin (BCG), a member of the *M. tuberculosis* complex, induces lipid droplet (LD) accumulation in murine macrophages in a TLR2-dependent manner, yet activation of TLR2 alone by the synthetic agonist Pam3CSK4 is insufficient to induce LD formation (15, 16). Similar results were obtained with BCG-derived ManLAM (15, 16). These observations left unresolved how BCG bacilli – and specifically ManLAM – promote LD biogenesis in macrophages. Because ManLAM is a relatively weak TLR2 agonist compared to bacterial lipoproteins (EC_50_ is in the µg/ml *vs* ng/ml range) (13, 17), it was important to exclude the possibility that the observed TLR2-stimulating activity of the ManLAM preparation resulted from trace contamination with highly active lipopeptides (18). To address this, purified ManLAM was treated with 1% H_2_O_2_, which inactivates lipoproteins by converting the N-terminal cysteine-thioether substructure required for TLR2 activation into sulfoxide derivatives (19). Untreated and H_2_O_2_-treated ManLAM were tested with a TLR2 reporter HEK293 cell line engineered to express alkaline phosphatase activity as a read-out of TLR2-dependent NF-κB activation (see *Methods*). H_2_O_2_ treatment had little effect on the TLR2 agonist activity of *M. tuberculosis* ManLAM (**Fig. 1A**) while completely abrogating TLR2 activation by the sentinel lipoprotein LrpG (**SI Fig. 1**). This result demonstrates that our ManLAM preparations are highly purified and express bona fide TLR2 agonism.

**Figure 1:**
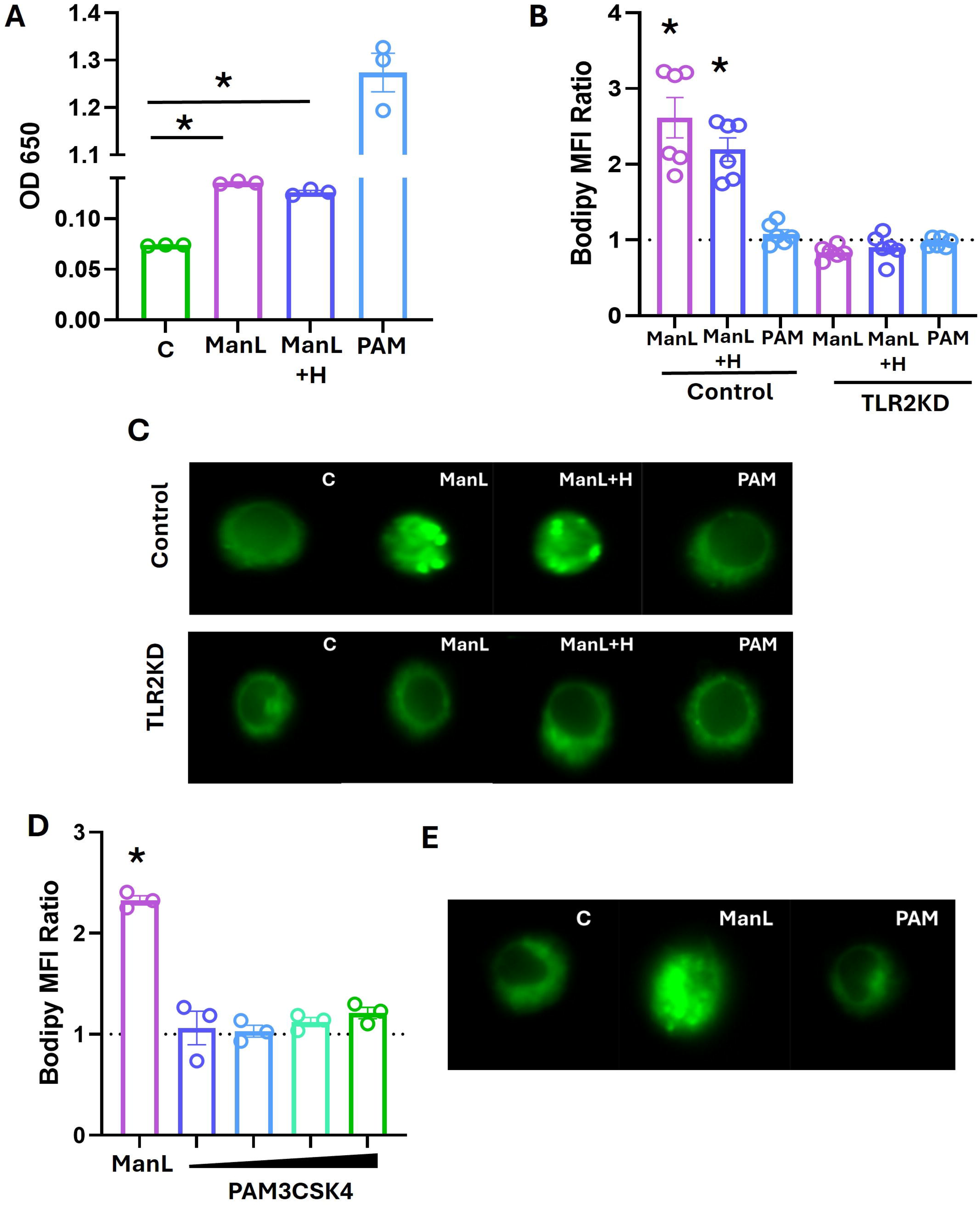
Involvement of TLR2 signaling in lipid droplet induction by *M. tuberculosis* ManLAM. **A.** *M. tuberculosis* ManLAM (500 ng/ml), H_2_O_2_-treated ManLAM (ManL+H; 500 ng/ml), Pam3CSK4 (PAM, synthetic TLR2 agonist; 500 ng/ml) and vehicle control (C) were tested for ability to induce NF-κB activation in HEK-TLR2 reporter cells. Reporter cells were treated for 24h and TLR2 activity was measured as a function of inducible secreted embryonic alkaline phosphatase in a colorimetric assay at 650nm. The Y-axis shows background-subtracted OD_650_ values. **B.** CRISPRi technology was used to generate TLR2 knock-down (KD) murine iBMDM. Control (non-targeting guide RNAs) and TLR2 KD clones were treated for 24h with 500ng/ml ManLAM preparations and Pam3CSK4. Abbreviations are as in panel A. Macrophages were stained with Bodipy 493/503 and antibodies to the macrophage marker F4/80. Data were obtained by imaging flow cytometry in triplicate and expressed as ratio of Bodipy mean fluorescence intensity (MFI) of treated vs. vehicle-treated cells. The horizontal dotted line marks the vehicle-treated cell reading. Comparison for statistical significance was between treated and vehicle-treated cells. **C**. Representative microscopy images (60x magnification) of vehicle-control (C) and treated iBMDM in the experiment shown in panel B. Fluorescence stain: Bodipy 493/503 (neutral lipid dye, green fluorescence). **D.** iBMDM were treated with ManLAM (500 ng/ml) and increasing doses of Pam3CSK4 (10, 50, 100, 500 ng/ml) for 24h. Data were generated and expressed as in panel B. **E.** Representative images (60x magnification) of the experiment shown in panel D. Description of the images as in panel C. *, *p* <0.05 by unpaired *t* test in all relevant panels.

Next, to investigate the mechanisms underlying ManLAM-induced LD accumulation in macrophages, we used immortalized murine bone marrow-derived macrophages (iBMDMs), which are well suited for genetic manipulations and closely recapitulate primary macrophage phenotypes (20). Gene knockdown in iBMDMs was achieved by CRISPR-interference (CRISPRi), which employs a nuclease-inactivated Cas9 (dCas9) and promoter-targeting guide RNAs (gRNAs) to block transcription of target genes. We generated TLR2 knockdown (KD) iBMDMs by stably expressing dCas9 and two different gRNAs targeting the *Tlr2* promoter (see *Methods*). Control cells expressed gRNAs against an irrelevant target gene (bacterial luciferase, *lux*A). Efficient TLR2 knockdown was confirmed by reduced *Tlr2* transcript levels measured by RT-qPCR (**Sl Fig. 2A**) and by impaired induction of TLR2-responsive genes *Il6* and *Tnfa* following stimulation with the TLR2 agonist Pam3CSK4 (**Sl Fig. 2B,C**). ManLAM and H_2_O_2_-treated ManLAM induced LD accumulation in control iBMDMs but failed to do so in TLR2 KD cells (**Fig. 1B**; representative imaging flow cytometry data in **Fig. 1C**). Moreover, in agreement with previous reports (16), stimulation of control iBMDMs with Pam3CSK4 alone failed to induce LD accumulation at any of the doses tested (**Fig. 1B-E**). The observed insufficiency of TLR2 signaling prompted us to investigate whether ManLAM engages other host receptors to drive macrophage LD accumulation.

**Figure 2:**
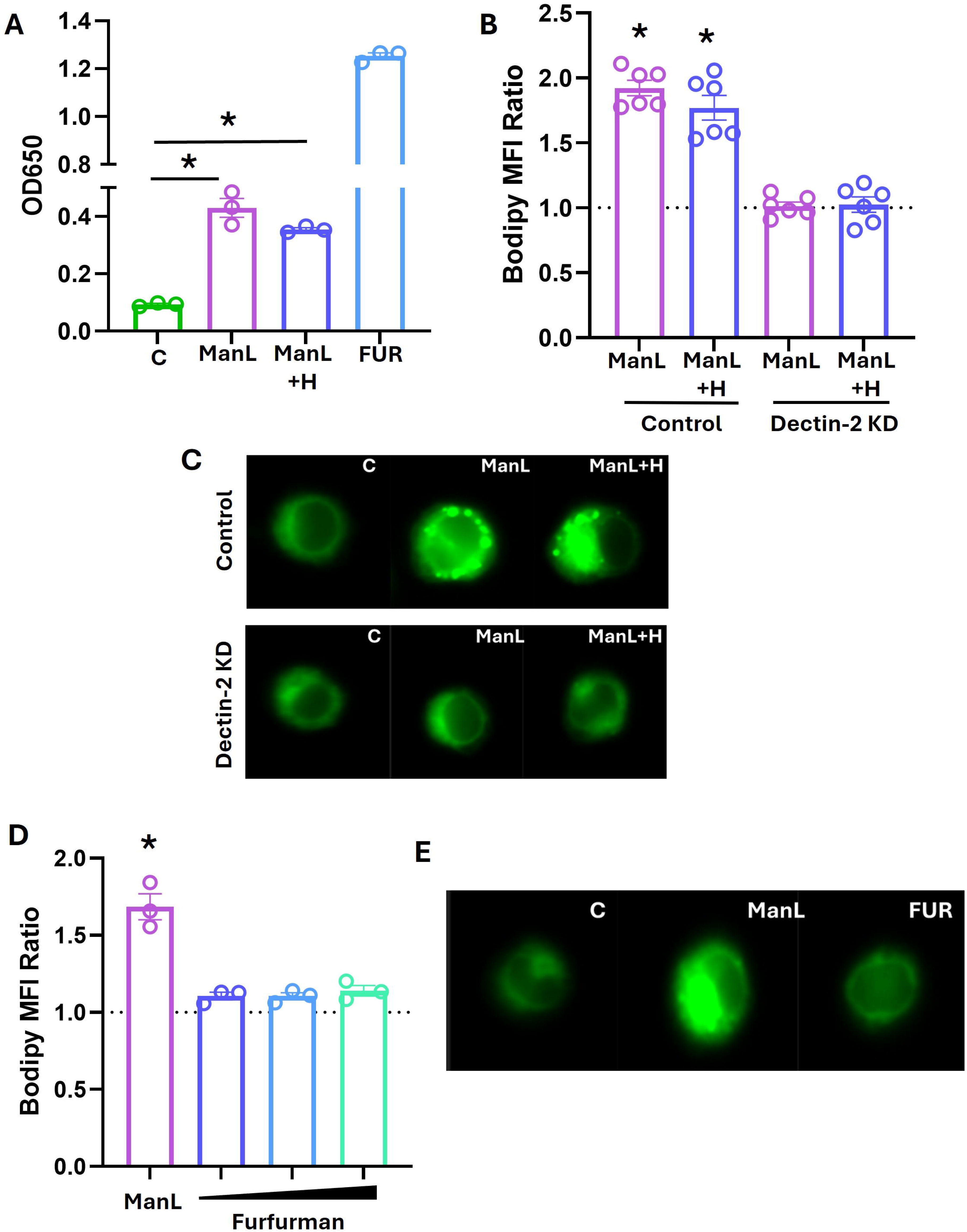
Involvement of Dectin-2 signaling in lipid droplet induction by *M. tuberculosis* ManLAM. **A.** *M. tuberculosis* ManLAM (500 ng/ml), H_2_O_2_ treated ManLAM (ManL+H, 500 ng/ml), furfurman (FUR; commercial Dectin-2 agonist; 10 µg/ml) and vehicle control (C) were tested were tested for ability to induce NF-κB activation in HEK-Dectin-2 reporter cells. Data were generated and analyzed as in Fig. 1A. **B.** Control and Dectin-2 knockdown (KD) iBMDM were treated 24h with 500ng/ml ManLAM preparations. Abbreviations are as in panel A. Imaging flow cytometry data were generated and expressed as in Fig. 1B. **C.** Representative microscopy images (60x magnification) of vehicle-control (C) and treated iBMDM in the experiment shown in panel B. Panel description as in Fig. 1C. **D.** iBMDM were treated with ManLAM (500 ng/ml) doses of furfurman (FUR; 2.5, 5 and 10 µg/ml). Imaging flow cytometry data were generated and expressed as in Fig. 1B. **E.** Representative microscopy images (60x magnification) of vehicle-control (C) and treated iBMDM in the experiment shown in panel D. Panel description as in Fig. 1C. *, *p* <0.05 by unpaired *t* test in all relevant panels.

Previous work has shown that *M. tuberculosis* ManLAM is also an agonist of Dectin-2 (11, 12), a surface CLR that recognizes microbial carbohydrates and has been studied primarily in the context of fungal infections (21). Consistent with these reports, *M. tuberculosis* ManLAM activated Dectin-2 signaling in a reporter HEK293 cell line engineered to express alkaline phosphatase activity as a read-out of Dectin-2-dependent NF-κB activation (see *Methods*) (**Fig. 2A**). H_2_O_2_ treatment had no significant effect on the Dectin-2 agonist activity of *M. tuberculosis* ManLAM (**Fig. 2A**). To assess whether Dectin-2 signaling contributes to ManLAM-induced LD accumulation, we generated Dectin-2 knockdown (KD) and control cells in iBMDM using the same strategy described for TLR2. Efficient Dectin-2 knockdown was validated by reduced *Dectin2* transcript levels measured by RT-qPCR (**SI Fig. 2D**) and by impaired induction of the downstream genes *Il6* and *Tnfa* following stimulation with the Dectin-2 agonist furfurman (**SI Fig. 2E,F**). When LD induction was assessed, both ManLAM and H_2_O_2-_treated ManLAM induced LD accumulation in control iBMDM but failed to do so in Dectin-2 KD cells (**Fig. 2B**; representative imaging flow cytometry data are in **Fig. 2C**). In contrast, knockdown of other CLRs, including Dectin-1 and Mincle, had no effect on ManLAM-induced LD accumulation (**Sl Fig. 3A**). Moreover, treating parental iBMDMs with increasing doses of furfurman up to 10 μg/ml failed to induce LD accumulation (**Fig. 2D**; representative imaging flow cytometry data are in **Fig. 2E**). Thus, as observed with TLR2, Dectin-2 activation is necessary but not sufficient to drive ManLAM-induced LD accumulation.

**Figure 3:**
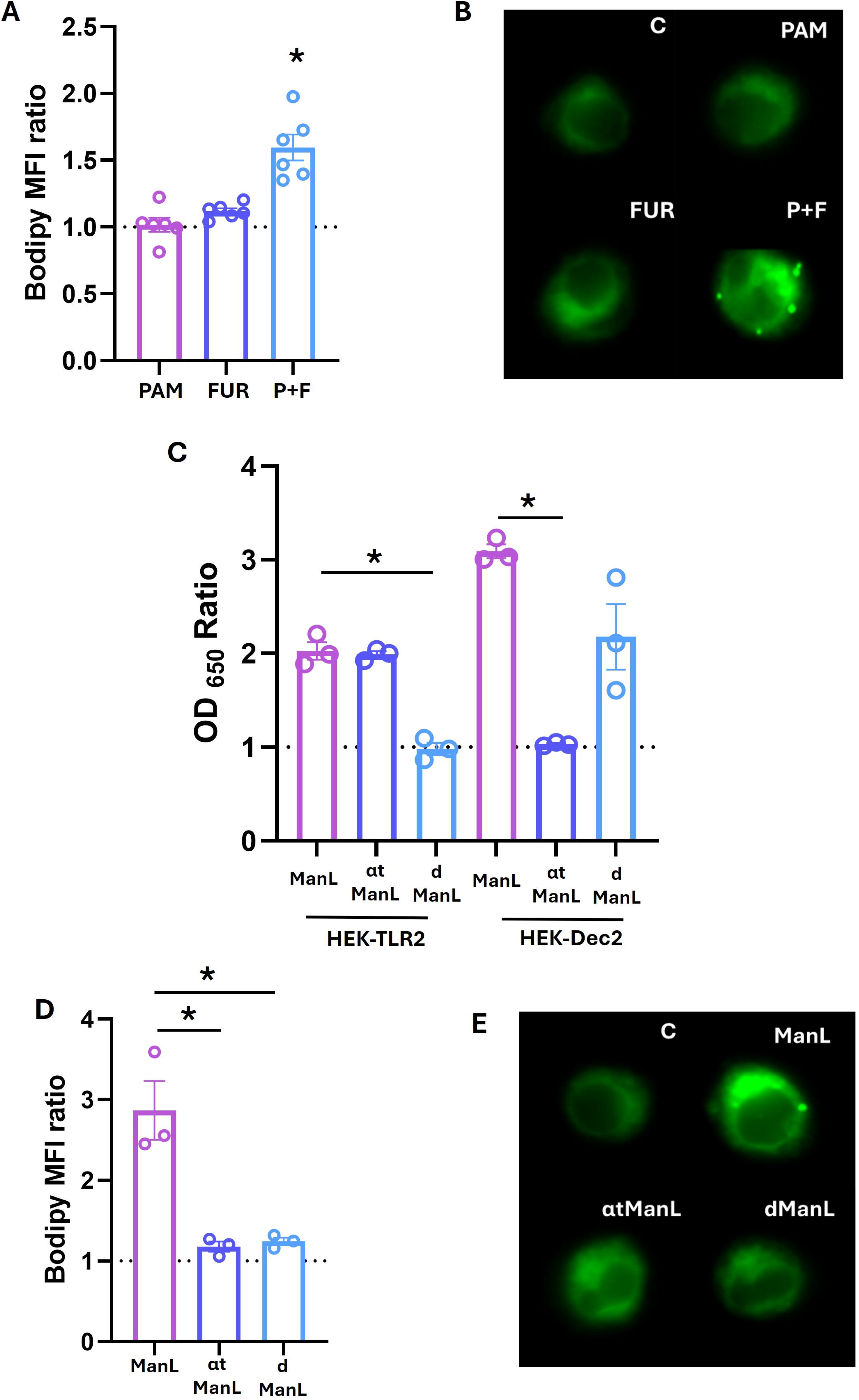
Involvement of *M. tuberculosis* ManLAM moieties in lipid droplet induction. **A.** iBMDM were treated with Pam3CSK4 (PAM; 500 ng/ml) and furfurman (FUR; 10 µg/ml), singly or in combination (P+R), and corresponding vehicle controls. Imaging flow cytometry data were generated and expressed as in Fig. 1B. **B.** Representative microscopy images (60x magnification) of vehicle-control (C) and treated iBMDM in the experiment shown in panel A. Panel description as in Fig. 1C. **C.** *M. tuberculosis* ManLAM, demannosylated ManLAM (αtManLAM), and deacylated ManLAM (dManLAM) (all H_2_O_2_ pre-treated, 500 ng/ml) and corresponding vehicle controls were tested for ability to induce NF-κB activation in HEK-TLR2 and HEK-Dectin 2 reporter cells. Treatment was for 24h as in preceding figures. Data are presented as ratio of OD_650_ values obtained with treated vs vehicle control cells. The horizontal dotted line marks the corresponding vehicle-control reading. **D.** ManLAM and derivatives, as in panel C, were used to treat iBMDM (all 500 ng/ml for 24h). Imaging flow cytometry data were generated and expressed as in Fig. 1B. **E.** Representative microscopy images (60x magnification) of vehicle-control (C) and treated iBMDM in the experiment shown in panel D. Panel description as in Fig. 1C. *, *p* <0.05 by unpaired *t* test in all relevant panels.

The data above strongly suggest that *M. tuberculosis* ManLAM engages both TLR2 and Dectin-2 signaling to induce LD accumulation in macrophages. To determine whether co-engagement of these two receptors is sufficient, we treated iBMDMs with the TLR2 agonist Pam3CSK4 and the Dectin-2 agonist furfurman, alone and in combination. While neither agonist induced LD accumulation when applied alone, they did so when used in combination (**Fig. 3A**; representative imaging flow cytometry data are in **Fig. 3B**). These results demonstrate that co-engagement of TLR2 and Dectin-2 is required to induce macrophage lipid droplet accumulation and raise the possibility that ManLAM functions as a single ligand that coordinates signaling through both receptors to drive macrophage lipid reprogramming.

We next asked whether specific ManLAM structural features are required for lipid droplet accumulation mediated by TLR2 and Dectin-2. Prior work showed that acylation of the phosphatidyl-myo-inositol anchor of mycobacterial lipoglycans is required for TLR2 activation (14, 22), whereas the mannose caps of ManLAM are required for recognition by Dectin-2 (11, 12). To selectively disrupt these interactions, we generated deacylated ManLAM by NaOH treatment (dManLAM, see *Methods*) and mannose-cap-deficient ManLAM by enzymatic removal with α-mannosidase (αtManLAM, see *Methods*). When tested in TLR2 and Dectin-2 reporter HEK293 cell lines, dManLAM failed to activate TLR2 while retaining Dectin-2 activation ability (**Fig. 3C**). In contrast, αtManLAM retained TLR2 activation but lost the ability to activate Dectin-2 (**Fig. 3C**). We next assessed the ability of these ManLAM derivatives to induce LD accumulation in iBMDMs. In contrast to intact ManLAM, neither dManLAM nor αtManLAM induced LD accumulation (**Fig. 3D**; representative imaging flow cytometry data are in **Fig. 3E**). These findings were recapitulated in human monocyte-derived macrophages (MDM) (**SI Fig. 3B**). Together, these results demonstrate that engagement of both TLR2 and Dectin-2 by ManLAM is required for macrophage LD accumulation, identify the structural moieties that mediate this dual receptor recognition, and establish its relevance in human macrophages.

### Structural integrity of ManLAM is required for full NF-**κ**B activation

Because ManLAM is also known to activate the transcription factor NF-κB (11), we next used NF-κB activation as a quantitative readout to compare signaling elicited by intact ManLAM with that induced by the structural derivatives used in the lipid droplet assays. NF-κB activation occurs downstream of multiple innate immune receptors, including TLRs and CLRs (23). Upon receptor engagement, a series of signaling events results in the translocation of NF-κB from the cytoplasm to the nucleus (schematic in **Fig. 4A**), where it induces expression of proinflammatory cytokines such as TNF-α and IL-1β (24). We therefore quantified NF-κB activation by measuring nuclear translocation of the RelA (p65) subunit of NF-κB and downstream cytokine gene expression in macrophages treated with ManLAM or its structural derivatives.

**Figure 4:**
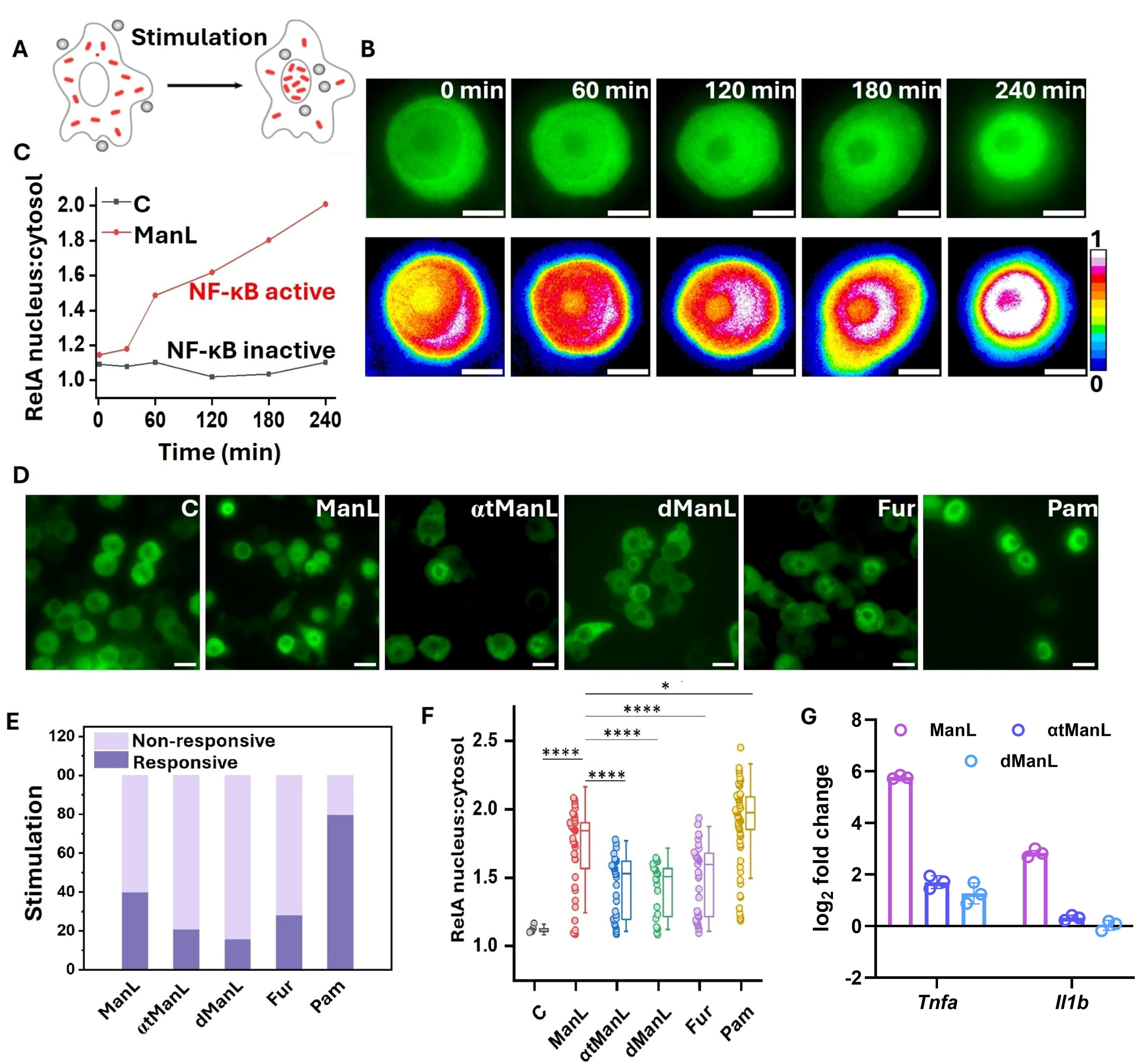
Ligand-dependent NF-κB RelA nuclear translocation dynamics in EGFP-RelA RAW264.7 macrophages. **A.** Schematic illustrating cytoplasm-to-nucleus translocation of NF-κB RelA (p65) upon macrophage cell stimulation. Grey circles, ManLAM and derivatives; red rods, RelA. **B.** (Top row) Time-lapse live-cell epifluorescence images showing EGFP-RelA translocation following ManLAM stimulation. (Bottom row) Same cell images shown with EGFP-RelA fluorescence intensity encoded in pseudo-color, as indicated by the color bar. Scale bars: 5 µm. **C.** Quantification of RelA nuclear translocation over time for the representative cell shown in panel B and for an unstimulated control cell, expressed as the fluorescence intensity ratio of EGFP-RelA in the nucleus relative to the cytoplasm. **D.** Representative epifluorescence images of fixed EGFP-RelA RAW264.7 macrophages stimulated with *M. tuberculosis* ManLAM, demannosylated ManLAM (αtManLAM), and deacylated ManLAM (dManLAM) or furfurman (FUR) for 240 min, or with Pam3CSK4 for 60 min, illustrating ligand-induced RelA nuclear translocation. Scale bars: 10 µm. **E.** Percentage of responsive and non-responsive cells under various ligand stimulation conditions, as indicated. Responsive cells were defined as those with a RelA nucleus-to-cytosol ratio greater than 1.089, a threshold set to achieve 95% of unstimulated cells being classified as non-responsive. Approximately 200 cells per condition were analyzed; the fraction of responsive cells was 40% for ManLAM, 21% for αtManLAM, 16% for dManLAM, 28% for furfurman (FUR), and 80% for Pam3CSK4. **F.** Single-cell scatter plots of NF-κB activation, expressed as the nucleus-to-cytosol EGFP-RelA fluorescence intensity ratio, in all responsive cells after 240 min ligand stimulation (for ManLAM and derivatives, and furfurman) and 60 min for Pam3CSK4. Each dot represents one cell. Box plots show the interquartile range and the median line; whiskers indicate ±1.5× SD. **G.** Expression levels of *Tnfa* and *Il1b* in RAW 264.7 cells treated with ManLAM, αtManLAM and dManLAM for 6 hours were measured by RTqPCR. Measurements were performed on triplicate samples and expressed as log_2_ fold change relative to the housekeeping actin gene. Statistical significance was assessed by one-way ANOVA: ***P < 0.001; **P < 0.01; *P < 0.05.

Nuclear translocation was calculated as the ratio of nuclear to cytoplasmic RelA fluorescence intensity in RAW 264.7 macrophages stably expressing green-fluorescence-protein-tagged RelA. Stimulation with ManLAM induced a gradual increase in nuclear RelA accumulation over time relative to unstimulated control cells (**Fig. 4B,C**). As expected, ManLAM-induced NF-κB activation was substantially less potent than that triggered by soluble Pam3CSK4, a benchmark agonist for robust TLR2-mediated activation.

ManLAM-induced RelA translocation exhibited substantial cell-to-cell variability, consistent with the intrinsic heterogeneity of immune cell activation (25). To quantify this response, we defined a threshold that classified 95% of unstimulated cells as nonresponsive. Using this criterion, the macrophage population segregated into two groups: a responsive subpopulation displaying robust nuclear RelA accumulation and a nonresponsive subpopulation remaining below the threshold (**SI Fig. 4**). Compared with intact ManLAM, stimulation with αtManLAM and dManLAM -- which selectively disrupt recognition by Dectin-2 and TLR2, respectively -- reduced both the fraction of responsive cells by approximately 50-60% (**Fig. 4D,E**) and the magnitude of RelA translocation within the responsive cell population (**Fig. 4F**), indicating attenuated NF-κB activation at both the population and single-cell levels. Since these structural modifications selectively disrupt Dectin-2 and TLR2 recognition, these findings indicate that full NF-κB activation by ManLAM requires coordinated engagement of both receptors. Consistent with this conclusion, intact ManLAM induced markedly greater expression of downstream cytokine genes in RAW 264.7 macrophages, including *Tnfa* and *Il1b*, with induction ∼25-fold and ∼6-fold higher, respectively, than that elicited by either structural derivative (**Fig. 4G**). Comparable results were obtained in iBMDM (**SI Fig. 5**), indicating similar responses across murine macrophage systems.

**Figure 5:**
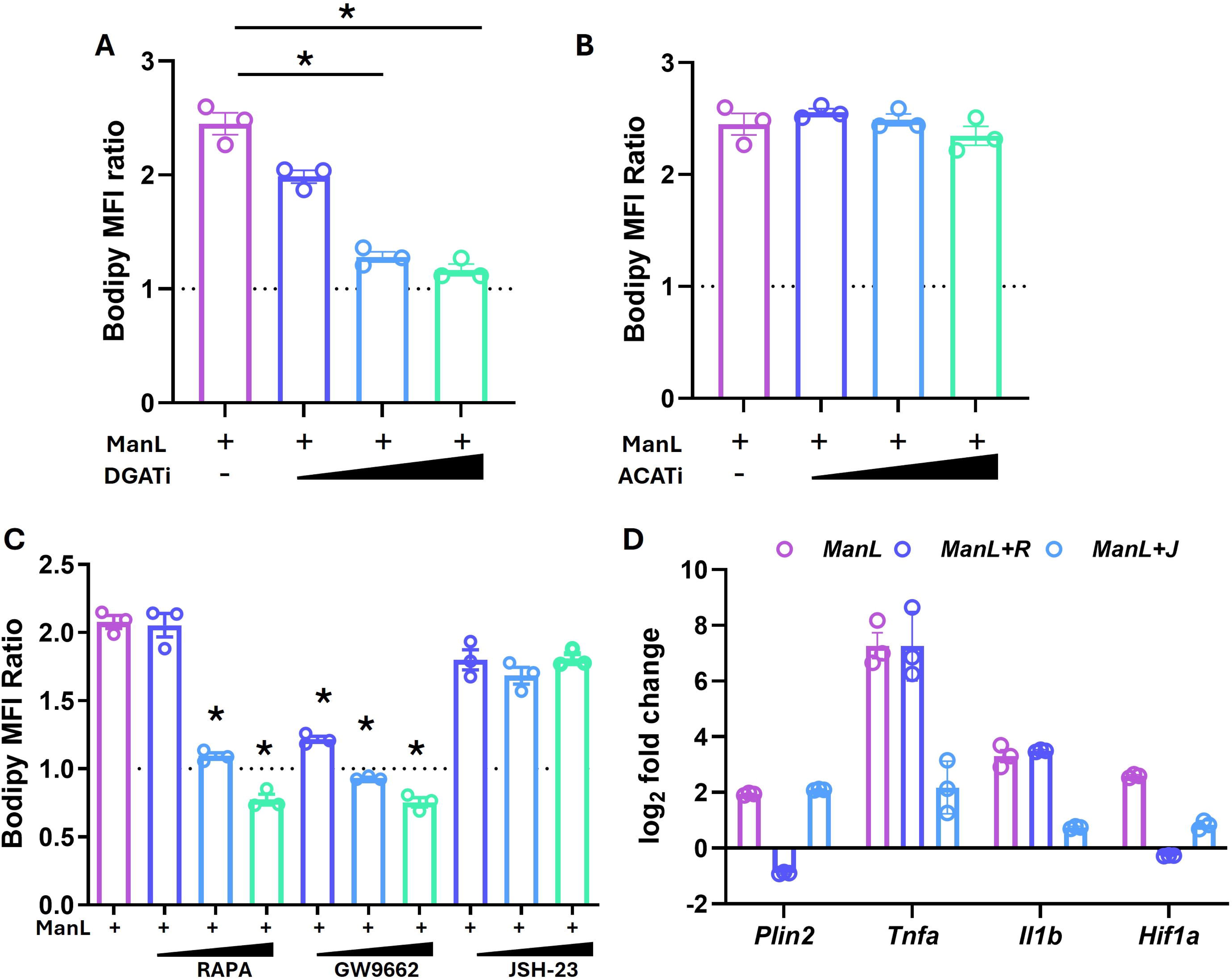
Effects of inhibitors of neutral lipid biosynthesis and NF-kB in ManLAM-treated macrophages. A,B. iBMDM were treated with ManLAM (500ng/ml) and vehicle-treated or treated with increasing doses of (A) the diacylglycerol transferase inhibitor, A922500 (DGAT-i; 60, 90, 120ng/ml) and (B) the acyl-coenzyme A:cholesterol acyltransferase inhibitor, CAS 615264-52-3 (ACAT-i; 5, 10, 15µg/ml). Imaging flow cytometry data for lipid droplet content were generated and expressed as in Fig. 1B. *, *p* <0.05 by unpaired *t* test in all relevant panels. **C.** iBMDM were treated with ManLAM (500ng/ml) and vehicle-treated or treated with increasing doses of the mTORC1 inhibitor rapamycin (0.2, 0.4, 0.8nM), PPARγ inhibitor GW9662 (0.5, 1, 2µM), and the NF-κB inhibitor JSH-23 (3.5, 7, 14µM). Imaging flow cytometry data for lipid droplet content were generated and expressed as in Fig. 1B. **D.** iBMDM were treated with ManLAM (500ng/ml) plus vehicle-control (ManL), the mTORC1 inhibitor rapamycin (ManL+R, 0.4nM), and the NF-κB inhibitor JSH-23 (ManL+J, 7µM) for 6 hours. Expression levels of *Plin2*, *Tnfa, Il1b, and Hif1a* were measured by RT qPCR in triplicate and expressed as log_2_ fold change, relative to the housekeeping actin gene. *, *p* <0.05 by unpaired *t* test in all relevant panels.

### ManLAM induces triglyceride-rich lipid droplets through an mTORC1-PPARγ pathway independent of NF-κB

We next investigated whether ManLAM-induced macrophage LD accumulation resembles that observed during live *M. tuberculosis* infection. To define the chemical nature of the ManLAM-induced neutral lipids, untreated and ManLAM-treated iBMDMs were exposed to chemical inhibitors of diacylglycerol acyltransferase (DGAT), which converts di- into tri-glycerides, or acyl-CoA:cholesterol acyltransferase (ACAT), which esterifies free cholesterol. Inhibition of DGAT, but not ACAT, reduced ManLAM-induced LD accumulation in a dose-dependent manner (**Fig. 5A,B**), indicating that ManLAM induces triglyceride-enriched LDs, consistent with our prior findings with *M. tuberculosis* infection (26).

We next examined the signaling pathways underlying ManLAM-induced LD formation. Previous work showed that LD accumulation during tuberculous mycobacterial infection requires the protein kinase Mechanistic Target of Rapamycin Complex 1 (mTORC1) and the downstream transcription factor peroxisome proliferator-activated receptor gamma (PPARγ) (16, 26, 27). Both are key regulators of cellular lipid metabolism (28–30). Inhibition of mTORC1 with rapamycin or of PPARγ with GW9662 markedly reduced ManLAM-induced LD accumulation in a dose-dependent manner (**Fig. 5C**), recapitulating findings in *M. tuberculosis* infection (16, 26, 27). In contrast, although ManLAM activates NF-κB, LD accumulation was unaffected by inhibition of RelA (p65) nuclear translocation by JSH-23 (**Fig. 5C**). Similar results were obtained with a second NF-κB inhibitor, QNZ (**SI Fig. 6A**), indicating that Man-LAM-induced LD formation is largely independent of NF-kB activity.

**Figure 6.**
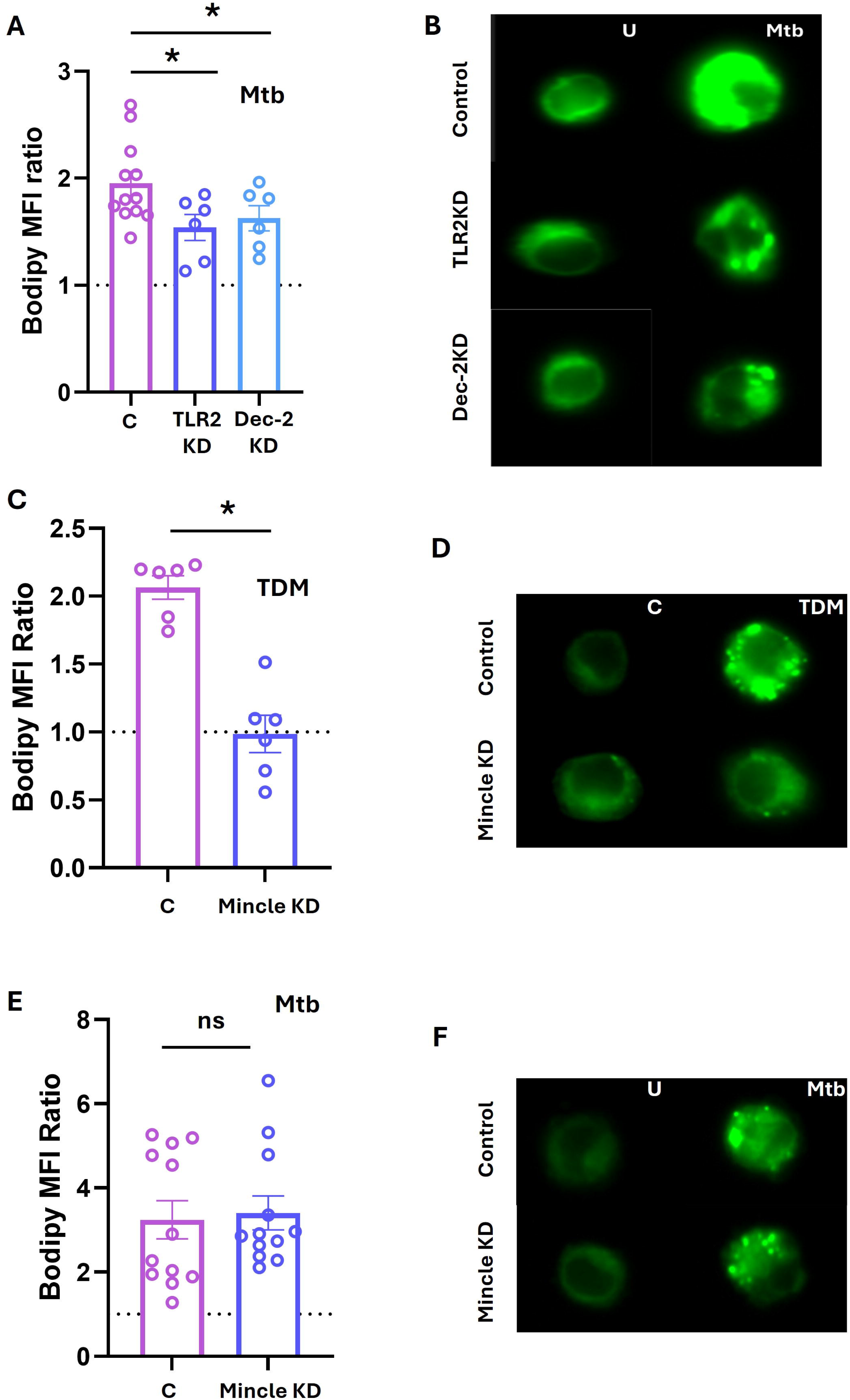
Effects on lipid droplet content of knocking down TLR2 and Dectin-2 in *M. tuberculosis*-infected macrophages and involvement of *M. tuberculosis* trehalose dimycolate (TDM) and Mincle in lipid droplet induction in vitro and during *M. tuberculosis* infection. **A.** Control (C, non-targeting guide RNAs), TLR2 knock-down (KD), and Dectin-2 (Dec-2) KD iBMDM clones were left uninfected or infected with *M. tuberculosis* (MOI= 15) for 48h in triplicate wells, washed, and stained with Bodipy 493/503 and antibodies to the macrophage marker F4/80 for imaging flow cytometry. Data were generated and expressed as in Fig. 1B. **B.** Representative microscopy images (60x magnification) of uninfected (U) and infected (Mtb) iBMDM in the experiment shown in panel A. Panel description as in Fig. 1C. **C.** Control (C, non-targeting guide RNAs) and Mincle knockdown (KD) iBMDM were treated 24h with 10µg/ml TDM preparations. Imaging flow cytometry data were generated and expressed as in Fig. 1B. **D.** Representative microscopy images (60x magnification) of vehicle-control (C) and treated iBMDM in the experiment shown in panel C. Panel description as in Fig. 1C. **E.** Control (C, non-targeting guide RNAs), and Mincle knockdown (KD) iBMDM clones were left uninfected or infected with *M. tuberculosis* and data were expressed as in panel A. **F.** Representative microscopy images (60x magnification) of uninfected (U) and infected (Mtb) iBMDM in the experiment shown in panel E. Panel description as in Fig. 1C.

To validate this pathway separation at the transcriptional level, we examined how inhibition of mTORC1 or NF-κB affected expression of representative target genes following ManLAM stimulation. Rapamycin selectively inhibited ManLAM-mediated upregulation of the PPARγ target gene *Plin2*, which encodes a lipid droplet-associated protein (31), but did not affect induction of the NF-κB-regulated genes *Tnfa* and *Il1b* (32) (**Fig. 5D**). Conversely, the NF-κB inhibitor JSH-23 suppressed ManLAM-mediated induction of *Tnfa* and *Il1b* but not *Plin2*. Both rapamycin and JSH-23 reduced ManLAM-induced upregulation of *Hif1a*, a gene controlled by both NF-κB and mTORC1 (33, 34). Comparable results were obtained with the NF-κB inhibitor QNZ (**Fig. S6B**). Together, these findings indicate that Man-LAM-induced LD accumulation proceeds through an mTORC1-PPARγ-dependent metabolic program that operates independently of NF-κB signaling, mirroring the pathway architecture observed during *M. tuberculosis* infection (26, 35).

### The ManLAM/TLR2/Dectin-2 axis contributes to macrophage lipid remodeling during *M. tuberculosis* infection

To assess the relevance of our findings with purified ManLAM to live *M. tuberculosis* infection, we tested whether TLR2 and Dectin-2 contribute to macrophage LD accumulation during infection. Macrophages infected with *M. tuberculosis* and subjected to knockdown of TLR2 or Dectin-2 exhibited an approximately 25% reduction in LD content compared with parental cells (**Fig. 6A**; representative imaging flow cytometry data in **Fig. 6B**), indicating that both receptors contribute to maximal LD accumulation during infection. Since ManLAM is the only known *M. tuberculosis*-derived agonist of Dectin-2 (11), these findings indicate that ManLAM represents a substantial, though not exclusive, contributor to the LD-inducing capacity of the pathogen.

We next asked whether other lipogenic mycobacterial envelope components identified in vitro contribute to LD accumulation during infection. Trehalose-6,6′-dimycolate (TDM), an abundant glycolipid and major virulence factor of *M. tuberculosis* (36, 37), also induced LD accumulation in iBMDM (**Fig. 6C,D**). This response required expression of its cognate receptor Mincle, linking TDM-induced LD accumulation to its established recognition by Mincle (38, 39) (**Fig. 6C,D ; Fig. S7** for Mincle knockdown efficiency data). However, in contrast to TLR2 and Dectin-2, knockdown of Mincle had no detectable effect on LD abundance in *M. tuberculosis*-infected macrophages (**Fig. 6E,F**). Thus, while ManLAM and TDM are both sufficient to induce LD accumulation in vitro, only the ManLAM-associated TLR2/Dectin-2 axis measurably contributes to lipid remodeling during live infection. Together, these findings support the conclusion that the ManLAM/TLR2/Dectin-2 axis captures a physiologically relevant component of the macrophage lipid remodeling response to *M. tuberculosis*.

## Discussion

Foam cells are the cellular hallmark of necrotizing TB lesions (26), yet how mycobacterial signals integrate to reprogram macrophage lipid metabolism remains unresolved. Here, we show that *M. tuberculosis* ManLAM acts as a key structural determinant of foam cell formation by coordinating engagement of TLR2 and Dectin-2. Distinct structural features of ManLAM --acylation and mannose capping -- selectively mediate activation of TLR2 and Dectin-2, respectively (11, 12, 14, 22), and are jointly required for triglyceride-rich lipid droplet accumulation. These findings explain earlier observations that TLR2 signaling is necessary but insufficient for lipid droplet formation. Lipid remodeling proceeds through an mTORC1-PPARγ axis that is largely independent of inflammatory signaling. Together, these results support a model in which ManLAM co-engages two macrophage surface receptors through distinct molecular moieties, reprogramming lipid metabolism and achieving full inflammatory activation via separable pathways (**Fig. 7**). Importantly, ManLAM-induced lipid remodeling mirrors that induced by *M. tuberculosis* infection in both neutral lipid composition and pathway dependence, whereas the mycobacterial glycolipid TDM, although sufficient to induce LD accumulation in vitro, does not measurably contribute through its cognate receptor pathway during infection. Together, these findings establish the ManLAM/TLR2/Dectin-2 axis as an infection-relevant driver of macrophage lipid metabolic reprogramming and demonstrate that reductionist ligand-based approaches can identify physiologically relevant host-pathogen signaling pathways.

**Figure 7:**
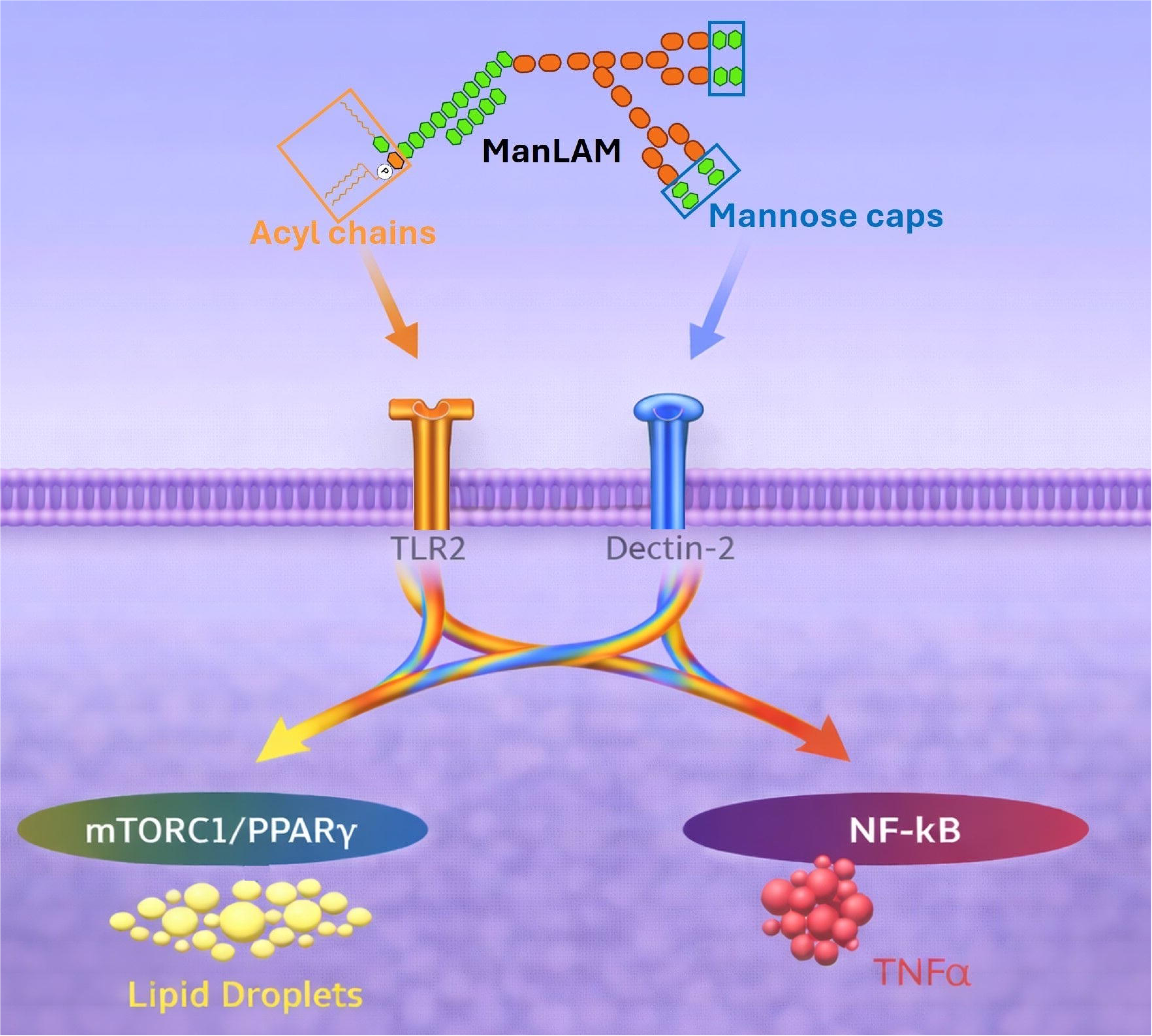
Dual receptor recognition of mannosylated lipoarabinomannan integrates macrophage signaling. Mannosylated lipoarabinomannan (ManLAM) engages TLR2 and Dectin-2 through distinct structural features. Lipid acyl groups interact with TLR2, whereas terminal mannose caps are recognized by Dectin-2. Signals from both receptors converge on shared intracellular pathways and diverge to activate two responses: mTORC1/PPARγ–dependent lipid droplet formation and NF-κB–dependent TNFα production. Illustration generated with AI-assisted tools and edited by the authors.

Activation of multiple receptors by distinct structural features within a single microbial ligand defines a mode of signal integration at the ligand level. In contrast, prevailing models of innate immune signaling emphasize receptor crosstalk, in which distinct microbial ligands engage separate receptors and signal integration occurs downstream of receptor activation – for example, cooperative signaling between CLRs and TLRs in fungal infections (40). Embedding ligands for multiple receptors within a single microbial component may confer an intrinsic advantage by promoting spatial proximity of engaged receptors. This idea is supported by prior studies showing that productive crosstalk between Dectin-1 and TLR2 depends on spatial coordination of their ligands, often requiring scaffolded presentation at nanometer-scale proximity (41, 42). Such spatial organization promotes receptor clustering, increases effective avidity, and converts weak interactions into productive signaling (43). By analogy, recognition of ManLAM by Dectin-2 and TLR2 may facilitate integration of parallel signaling cascades, enabling synergistic signaling even when individual receptor-ligand interactions are of low affinity. Ligand-level coordination may therefore represent a strategy by which microbes with complex, lipoglycan-rich cell envelopes -- including mycobacteria and fungi – integrate host sensing pathways.

ManLAM-driven lipogenic and inflammatory programs in macrophages appear to be functionally separable. This is consistent with established signaling biology, as TLR2 and Dectin-2, in addition to activating NF-kB (23), can independently engage the mTORC1-PPARγ axis through PI3K/Akt signaling downstream of MYD88 or Syk, respectively (44, 45). At the same time, the synergistic effects of TLR2 and Dectin-2 co-engagement on both lipogenic and inflammatory outputs suggest convergence at shared regulatory nodes. One candidate is the PI3K/Akt axis, which lies upstream of both NF-kB and mTORC1–PPARγ signaling (46). ManLAM may therefore promote foam cell formation by activating parallel signaling pathways that convergence at common control points while preserving functional independence of downstream outputs.

ManLAM-induced lipid remodeling closely mirrors that induced by *M. tuberculosis* infection in both neutral lipid composition and pathway dependence (26), and TLR2 or Dectin-2 deficiency reduces LD accumulation during infection. Since ManLAM is the only known *M. tuberculosis*-derived agonist of Dectin-2 (11), these findings support the conclusion that the ManLAM/TLR2/Dectin-2 axis captures a physiologically relevant component of the macrophage lipid remodeling response to *M. tuberculosis*. However, the partial reduction in LD accumulation observed in TLR2- or Dectin-2-deficient macrophages indicates that ManLAM is not the sole driver of this response in vivo and that additional microbial or host-derived signals likely contribute to foam cell formation during tuberculosis. Indeed, additional mycobacterial ligands and host-derived signals can induce LD accumulation (26, 38, 47–49). Our findings with the TDM/Mincle axis indicate that not all such pathways may contribute during infection, or that they may instead do so within distinct microenvironmental contexts or at later stages of lesion development. Together, these results support a hierarchical model in which ManLAM acts as a central driver of macrophage lipid remodeling during *M. tuberculosis* infection, whereas additional microbial and host-derived signals shape the magnitude, persistence, and progression of lipid-laden macrophages toward foam cell formation.

By identifying mTORC1 and PPARγ as downstream effectors of dual receptor engagement, we define key nodal control points within the lipid metabolic program that may be targeted pharmacologically to disrupt pathogenic foam cell formation. Because these regulators lie downstream of receptor engagement, their modulation may selectively constrain lipid remodeling while preserving upstream innate immune recognition. Thus, elucidating how ligand architecture encodes receptor integration not only advances our understanding of host–pathogen signaling logic but also highlights mechanistically grounded opportunities for host-directed therapy.

## Materials and Methods

### Mycobacterium tuberculosis components

Mannosylated lipoarabinomannan (ManLAM) was extracted from *M. tuberculosis* Erdman and purified by a previously described method (50, 51). Deacylated ManLAM (dManLAM) was prepared as described previously (50). Briefly, 200 µg of ManLAM was incubated in 200 µl of NaOH 0.1 M for 2 h at 37°C. After neutralization with 200 µl of HCl 0.1 M, the reaction products were dialyzed against water. ManLAM devoid of mannose caps (αtManLAM) was prepared as described previously (52). Briefly, 200 µg of ManLAM was incubated for 6 h at 37°C in 30 µl of α-mannosidase solution (2 mg/ml, 0.1 M sodium acetate buffer, pH 4.5, 1 mM ZnSO_4_) followed by a second addition of 50 µl of the same enzyme solution for overnight incubation at 37°C. The reaction products were dialyzed against 50 mM NH_4_CO_3_, pH 7.6. Elimination of α-mannosidase was achieved by denaturation (2 min at 110°C) followed by overnight tryptic digestion with trypsin/α-mannosidase (2% by weight) at 37°C. αtManLAM was recovered after dialysis against water. Removal of mannose caps was controlled by capillary electrophoresis as described (52). dManLAM and αtManLAM were suspended in 1x phosphate buffer saline (PBS) pH 7.4 and stored at -20°C. To inactivate putative lipoprotein contaminating ManLAM and its derivatives, samples were incubated with 1% H_2_O_2_ for 3h at 37°C, a treatment that converts the N-terminal cysteine-thioether substructure of lipoproteins into TLR-2-inactive sulfoxide derivatives (19). At the end of the incubation period, H_2_O_2_ was evaporated under N_2_ gas and the resulting pellets were reconstituted in distilled H_2_O.

An analog of *M. tuberculosis* trehalose-6,6′-dimycolate (TDM), GlcC14C18, was synthesized chemically, according to methods described in Decout *et al* (53).

### Cell lines

Murine bone marrow marrow-derived macrophages obtained from C57BL/6 mice and immortalized with J2 retrovirus (iBMDMs) (54) were kindly provided by Dr. Phillip West (Jackson Laboratory, Bar Harbor, ME). iBMDMs and RAW 264.7 cells (ATCC TIB-71) were cultured in Dulbecco’s modified Eagle’s medium (DMEM) supplemented with heat-inactivated 10% FBS, 1x penicillin-streptomycin solution (100 IU/mL penicillin and 100 μg/ml Streptomycin; Corning, Manassas, VA) and 20 mM HEPES (Millipore, Billerica, MA) and incubated at 37°C in a humidified atmosphere consisting of 5% CO_2_. RAW 264.7 cells carrying an enhanced green fluorescence protein (EGFP)-RelA reporter (25) were cultured in DMEM containing 10% FBS, 2 mM l-glutamine, 1x penicillin-streptomycin solution (Corning, Manassas, VA), and 10 mM HEPES (Millipore, Billerica, MA) at 37°C and 5% CO_2_. The HEK-Blue mDectin-2 and HEK-Blue hTLR2 (InvivoGen, San Diego, CA) are derivatives of HEK293 cells that stably express the murine Dectin-2 and the human TLR2 genes, respectively, along with a NF-κB-inducible reporter system (secreted alkaline phosphatase). These cells were maintained in DMEM containing 10% FBS, 4.5 g/l glucose, 2 mM L-glutamine, 1x penicillin-streptomycin solution, 100LJμg/ml zeocin, 200 μg/ml hygromycin, 10 μg/ml blasticidin, 1 μg/ml puromycin, and 1x TLR and CLR Selection solution (all from InvivoGen, San Diego, CA). Reporter cells were cultured for two passages without selection before use.

### Gene knock-down (KD) in iBMDMs

iBMDMs were transduced with lentivirus carrying nuclease-dead FLAG-tagged Cas9 (dCas9) and selected for using blasticin (5-10 μg/ml) (InvivoGen, San Diego, CA). The Lenti-dCas9-KRAB-blast plasmid was a gift from Gary Hon (U. Texas Southwestern) (Addgene plasmid # 89567; http://n2t.net/addgene:89567) (55). For each gene of interest, 5-6 small guide RNAs (sgRNAs) were selected using the CRISPick tool (Broad Institute, https://portals.broadinstitute.org/gppx/crispick/public) (56, 57). sgRNAs were cloned into the pLentiGuide-Puro backbone, which was a gift from Paul Khavari (Stanford U.) (Addgene plasmid # 117986; http://n2t.net/addgene:117986). dCas9-containing cells were transduced with lentivirus carrying gene-specific sgRNAs and selected for by using puromycin (5-10 μg/ml) (InvivoGen, San Diego, CA). Non-targeting sgRNAs (targeting *luxA*) were used as negative controls: *lux1*: GGCAATGAAACGCTACGCTC, *lux2*: ATAAAGAGCGCGCCCAACAC.

Knockdown cells were validated by RT-qPCR, and the cell lines carrying the two gene-specific sgRNAs with the highest knockdown efficiency were used for subsequent experiments. The gene-specific sgRNAs used in this study were: TLR2-1: TGGGTGTCCCTCTTCCTGCA, TLR2-2: AAGCTGATCCGCCCGGCTGG; Dectin-2-1: GGACCTGGCTTCTGTCAAAG, Dectin-2-2: TGAGTTAAATGCCACAGAGC; Mincle-1: TGATAGAAAAGCACTTACTG, Mincle-2 CAGTAAGTGCTTTTCTATCA. For both steps of transduction, lentivirus was produced using HEK293T-Lenti-X cells (Takara) and psPAX2 (Addgene plasmid #12260; http://n2t.net/addgene:12260) and pMD2.G/VSV-G (Addgene plasmid #12259; http://n2t.net/addgene:12259) packaging plasmids, which were gifts from Didier Trono (EPFL, Swiss Federal Technology Institute of Lausanne).

### Human monocyte-derived macrophages

Human leukopaks were obtained from the New York Blood Center (Long Island City, NY, USA). Peripheral blood mononuclear cells (PBMC) were prepared by Ficoll density gradient centrifugation (Ficoll-Paque, GE Healthcare, Uppsala, Sweden) and human monocyte-derived macrophages (hMDM) were cultivated as described in previous work (26). Treatments were performed on day 7 of culture.

### M. tuberculosis infection

*M. tuberculosis* H_37_Rv strain mc^2^6206 (Δ*panCD* Δ*leuCD*) was kindly provided by Dr. William Jacobs Jr. (Albert Einstein College of Medicine) (58). The strain was confirmed positive for phtiocerol dimycocerosates (PDIM) both initially by the Jacobs lab via thin layer chromatography of lipid extracts and during its successive culturing by using vancomycin susceptibility (VAN10) assays (59). To prevent PDIM loss, bacteria were cultured in media containing propionate (59). Prior to infection experiments, the strain was grown in Middlebrook 7H9 supplemented with 10% OADC, 0.5% glycerol, 0.05% Tween-80, 24 ug/ml pantothenate, 50 ug/ml leucine, and 0.1 mM propionate. Macrophage infections were performed as previously described (60, 61). Briefly, *M. tuberculosis* cultures were grown to an OD_600_ of 0.6-1.0, washed three times with PBS, and centrifuged at 500 x *g* for 5 min to remove clumps. Bacteria were diluted in DMEM plus 10% horse serum and added to iBMDMs plated at 3-4x10^5^ cells/well in a 12-well dish in iBMDM media containing 24 ug/ml pantothenate and 50 ug/ml leucine. A diluted bacterial inoculum was added to cultured iBMDM (MOI = 15), and cells were centrifuged at 1,000 x *g* for 10 min to synchronize the infection and resuspended in fresh iBMDM media containing 24 ug/ml pantothenate and 50 ug/ml leucine; culture media was changed daily. After 48 h, iBMDMs were detached from tissue culture plates by resuspending in 1 x PBS plus 4mM EDTA by gentle scraping. An equal volume of 8% paraformaldehyde was added to fix and inactivate the samples. Cells were washed twice with 1 x PBS and processed for LD formation as described below.

### Chemical reagents and antibodies

TLR2 ligand Pam3CSK4 and Dectin-2 ligand furfurman were purchased from Invivogen (San Diego, CA). Inhibitors of diacylglycerol transferase (A922500) and acyl-coenzyme A:cholesterol acyltransferase (CAS 615264-52-3), both from Santa Cruz Biotechnology, Dallas, TX, were stored as per manufacturer’s instructions. Inhibitors of Mechanistic Target of Rapamycin Complex 1 (Rapamycin), peroxisome proliferator-activated receptor gamma (GW9662), NF-κB (JSH-23 and QNZ) were all purchased from Selleckchem, Houston, TX. Inhibitor doses were selected at and around published EC_50_ data and toxicity profiles in macrophages (26, 62). Only inhibitor doses yielding >85% cell viability were utilized by using trypan blue (Cytiva, Marlborough, MA) and counting with an automated cell counter (LUNA-FX7, Logos Biosystem, Annandale, VA). Mouse Fc receptor blocking solution, anti-mouse CD16/CD32, BV421 rat anti-mouse F4/80, BV421 mouse anti-human CD11c (BD Biosciences, San Diego, CA) and Bodipy 493/503 (Life Technologies, Carlsbad, CA) were used at the dilutions suggested by the manufacturers and as per previous work (26).

### TLR2 and Dectin-2 reporter cell experiments

The HEK-Blue reporter cell experiments were performed following manufacturer’s protocols (13), with minor modifications. For the HEK-Blue hTLR2 reporter assay, 20 µl of ManLAM, dManLAM, αtManLAM, and Pam3CSK4 (TLR2 ligand), all suspended in 1x PBS, were added to each well in 96-well plates (in triplicate) to obtain a final concentration of 5 µg/ml of each reagent. HEK-Blue hTLR2 cells were harvested, washed in 1x PBS, and resuspended in Quanti-Blue media (InvivoGen, San Diego, CA) at a final density of 280,000 cells/ml. 180 µl of cell suspension was added to each well and incubated for 24 h at 37°C in a humidified atmosphere containing 5% CO_2._ For the HEK-Blue mDectin-2 reporter assay, 10 µl of ManLAM, dManLAM, and αtManLAM, and furfurman (Dectin-2 ligand) were suspended in 1x PBS and added to each well in 96-well plates (in triplicate) to obtain a final concentration of 10 µg/ml for ManLAM and derivatives and 200 µg/ml for furfurman. The plates were dried at 37 °C until the solvent evaporated (∼3 h). HEK-Blue mDectin-2 cells were harvested, washed in 1x PBS, and resuspended in Quanti-Blue media at a final density of 280,000 cells/ml. 200 µl of cell suspension was added to each well and incubated for 24 h at 37°C in a humidified atmosphere containing 5% CO_2._ For both reporter cell lines, the colorimetric reading was taken at 650 nm (OD_650_). The data were expressed as background-subtracted OD_650_ or OD_650_ ratio between treated cells and vehicle control cells.

### Lipid droplet measurements

One million murine iBMDM cells or one million human PBMCs in 1 ml culture medium were added to each well of 12-well tissue culture plates. Incubation was 2 h for murine cells and 7 days at 37°C in a humidified atmosphere containing 5% CO_2_. *M. tuberculosis* components and control agonists were diluted in the same medium and 1 ml of each suspension or solution was added to each well to obtain final concentrations of 500 ng/ml for ManLAM, ManLAM derivatives, and Pam3CSK4, and 10 µg/ml for furfurman. Treated and vehicle control cells were incubated for 24 h at 37°C in a humidified atmosphere containing 5% CO_2._ When appropriate, chemical inhibitors were added at the indicated doses in a 10 µl solution, concurrently with the treatments above.

After incubation, cells were subjected to imaging flow cytometry as described (26). Briefly, medium was discarded and 1 ml 1x PBS was added to each well. Macrophages were detached from culture plates by gentle scraping, pelleted by centrifugation at 350g for 5 min, and fixed with 4% paraformaldehyde in 1x PBS for 45 min at RT. Cells were washed with 1x PBS containing 0.1% bovine serum albumin (PBS-BSA), resuspended in 50 μl of PBS-BSA containing 5 μl of mouse Fc-receptor blocking solution, and incubated at RT for 7 min. After incubation, 50 μl of PBS-BSA containing 5 μl of mouse F4/80-BV421 or human CD11c-BV421 antibody were added to each tube, and samples were incubated for 30 min at 4°C. After washing with PBS-BSA, cells were stained with 0.3 μg/ml Bodipy 493/503 in 1x PBS for 15 min. For each condition, data were acquired from 5,000-10,000 F4/80-positive cells utilizing an ImageStreamXMark II imaging flow cytometer (Amnis Corporation, Seattle, WA), using 60x magnification. Image data were analyzed by IDEAS software version 6.0 (Amnis Corporation, Seattle, WA) after applying a compensation matrix and selecting the region of interest (lipid droplets) with the Spot Mask tool. Mean fluorescence intensity (MFI) and spots per cell were extracted and expressed as ratio of treated vs its vehicle control cells.

### Real Time PCR analysis

RNA was isolated using RNeasy Mini Kit (Qiagen, Valencia, CA, USA). Isolated RNA was subjected to cDNA synthesis using the SuperScriptTM First-Strand Synthesis System (Bio-Rad, Hercules, CA). Real time PCR was performed in a QuantStudio 7 Flex Real-Time time PCR machine (Thermo Fisher Scientific, Waltham, MA, USA) using the ABsolute QPCR SYBER Green Mix (ABgene, Rochester, NY, USA), according to the standard ABgene protocol. The sequence of the primers used for real time PCR are listed in **SI Table 1**. The internal control gene β-Actin was amplified simultaneously in a separate reaction. The threshold cycle number (CT) was determined by qPCR for triplicate reactions, and the mean CT of the triplicate reactions was calculated. Levels of expression of the genes of interest were normalized to β-Actin using the formula 2^−ΔDCT^, where –ΔDCT = ΔCT (treated) – ΔCT (control), and ΔCT is the CT of the target gene subtracted from the CT of the housekeeping gene β-Actin.

### Measurements of RelA nuclear translocation (NF-κB activation)

For the Rel-A nuclear translocation assay, RAW264.7 cells carrying a EGFP-RelA reporter gene fusion were plated at 4□×□10^5^ cells per well in a 96-well glass-bottom plate overnight and then serum starved for 3 h. Cells were stimulated with the same treatments used for lipid accumulation experiments, and incubated for 0.5, 1.0, 1.5 and 2.0 h at 37°C. For fixed cell imaging, cells were fixed after stimulation with 2% PFA on ice for 5 min. Nuclei were counterstained with 2 μg/mL Hoechst stain at RT for 15 min. Fluorescence images were acquired using a Nikon Eclipse Ti microscope equipped with an Andor iXon3 EMCCD camera and a Nikon Plan Apo 40□×□/0.95 objective. For each sample, 20 images were taken at different locations. Images were analyzed using ImageJ following a previously reported method (25).

### Statistical analysis

All values are presented as means ± standard deviation (SD) for technical triplicates. Comparisons between two groups were performed using a two-tailed Student’s t-test in the Prism graphpad software or by one-way ANOVA. The criterion for statistical significance was p < 0.05.

## Acknowledgements

We thank John Chan and Karl Drlica for critical comments on the manuscript, and staff of the Flow Cytometry and Immunology Core Laboratory at Rutgers New Jersey Medical School for assistance with imaging flow cytometry. This work was funded in part by AHA fellowship 24PRE1198252 (J.R.), NIH grants R01HL149450, R01AI158911, NCATS UM1TR004789 (M.L.G.), R35GM124918 (Y. Y.), NIH DP2AI154429 (S.B.), and a grant from the Fondation pour la Recherche Médicale Equipes FRM DEQ20180339208 (J.N.).

## Author contributions

D.N., M.G., S.B., J.N., and Y.Y. contributed to the experimental design; D.N., J.R., O.V., S.B., J.C., and J.N. performed experiments and analyzed data; D.N. and M.G. wrote the manuscript. All authors reviewed and edited the manuscript.

---

**Figure S1.**
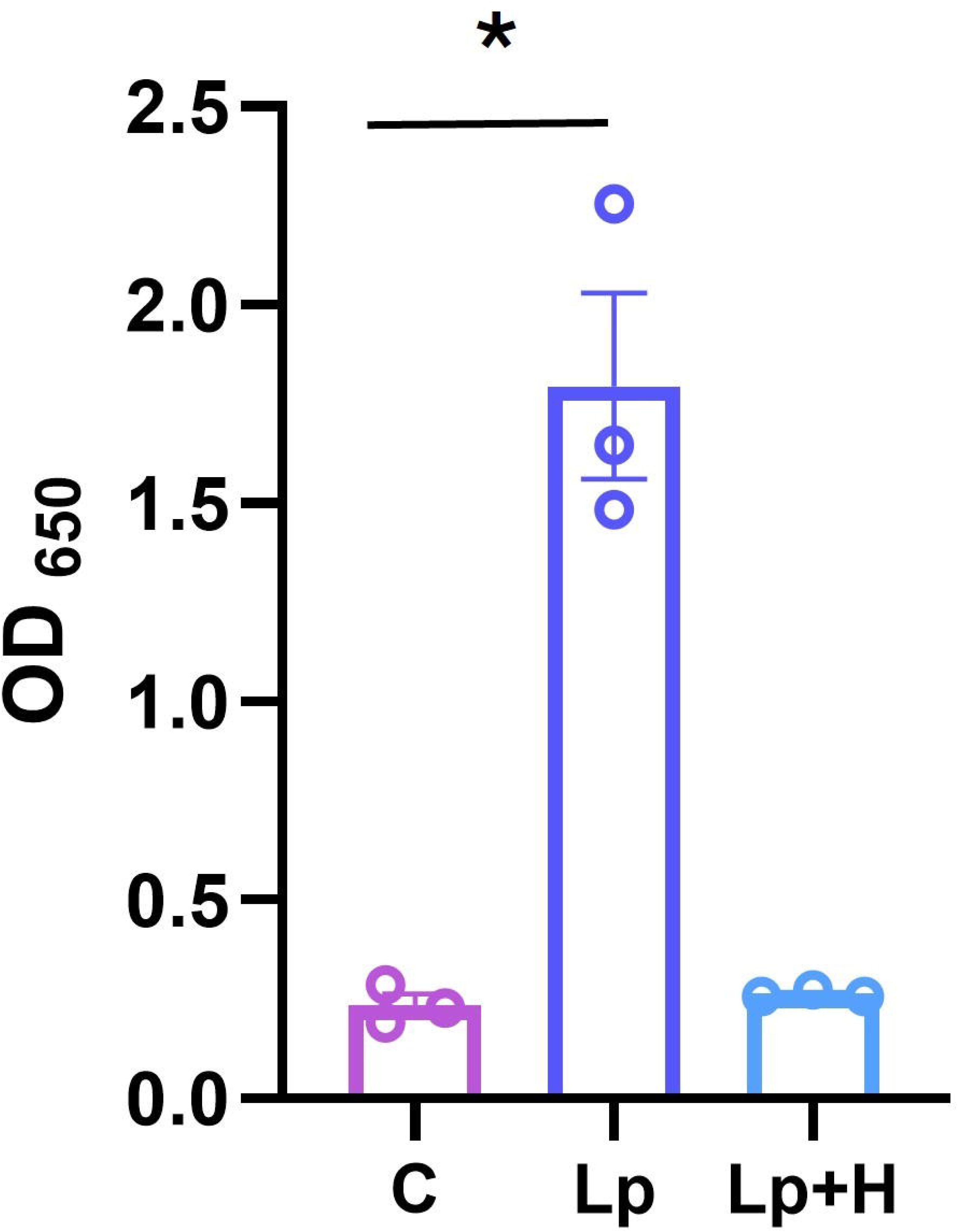

**Figure S2.**
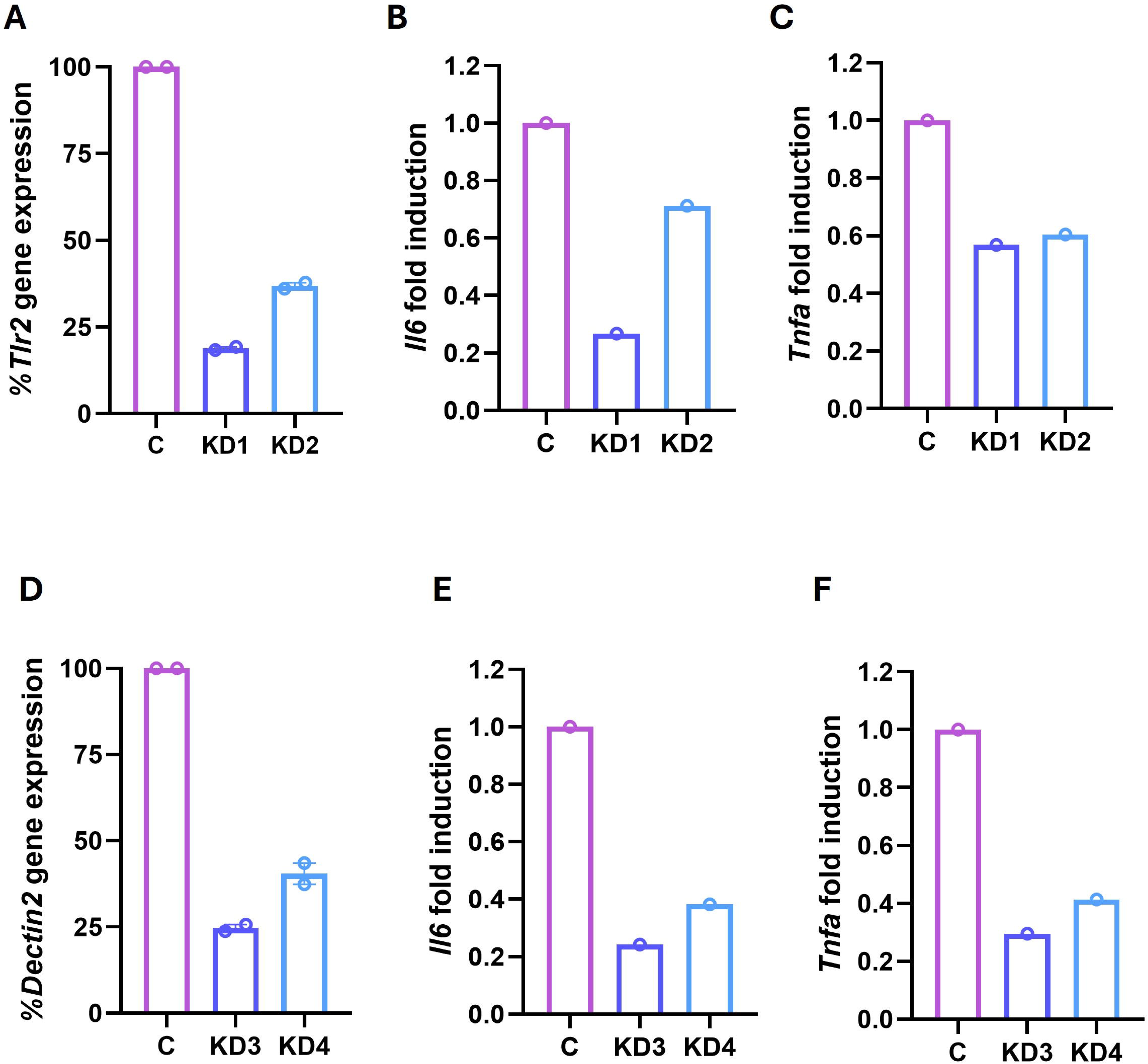

**Figure S3.**
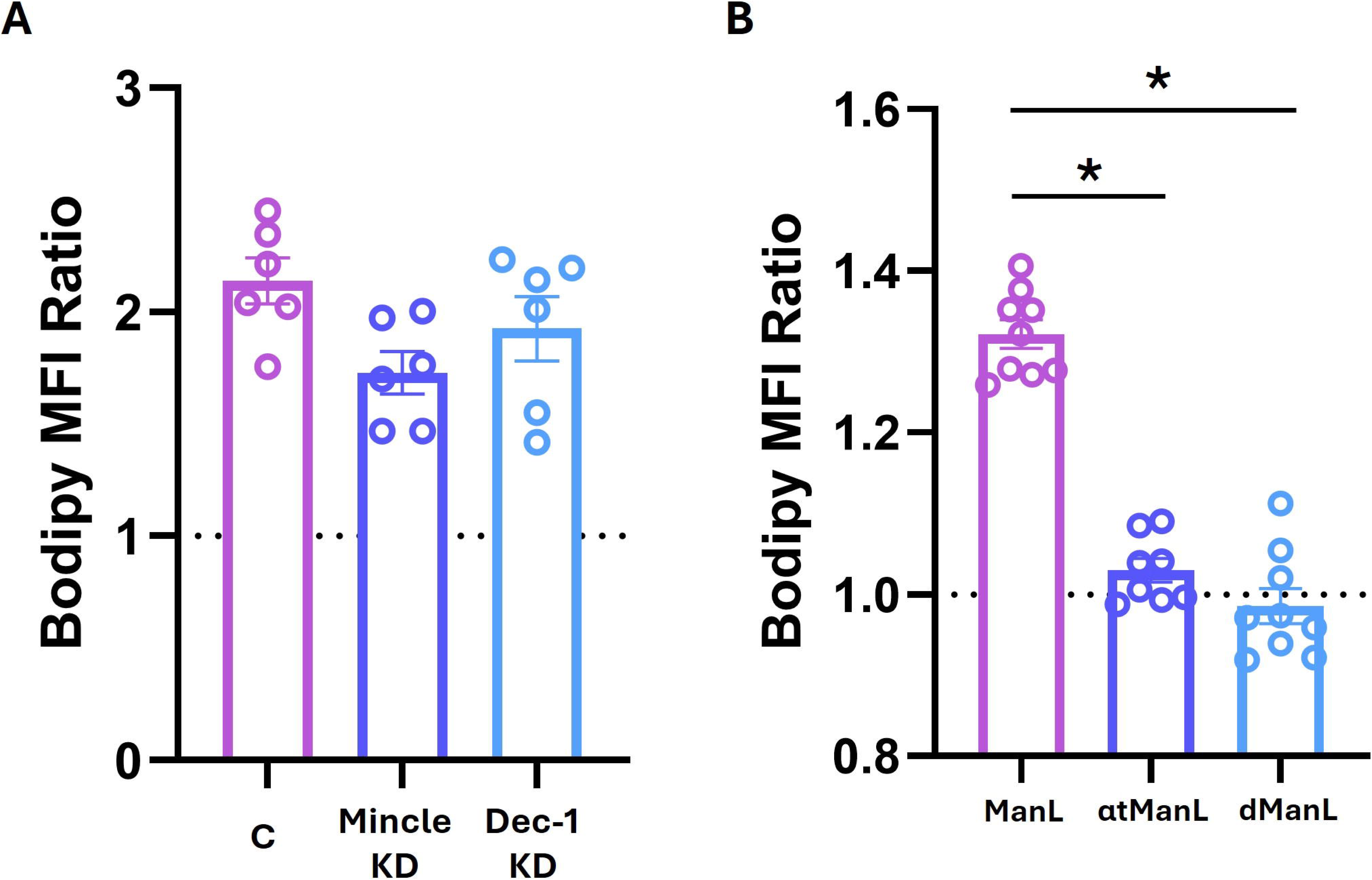

**Figure S4.**
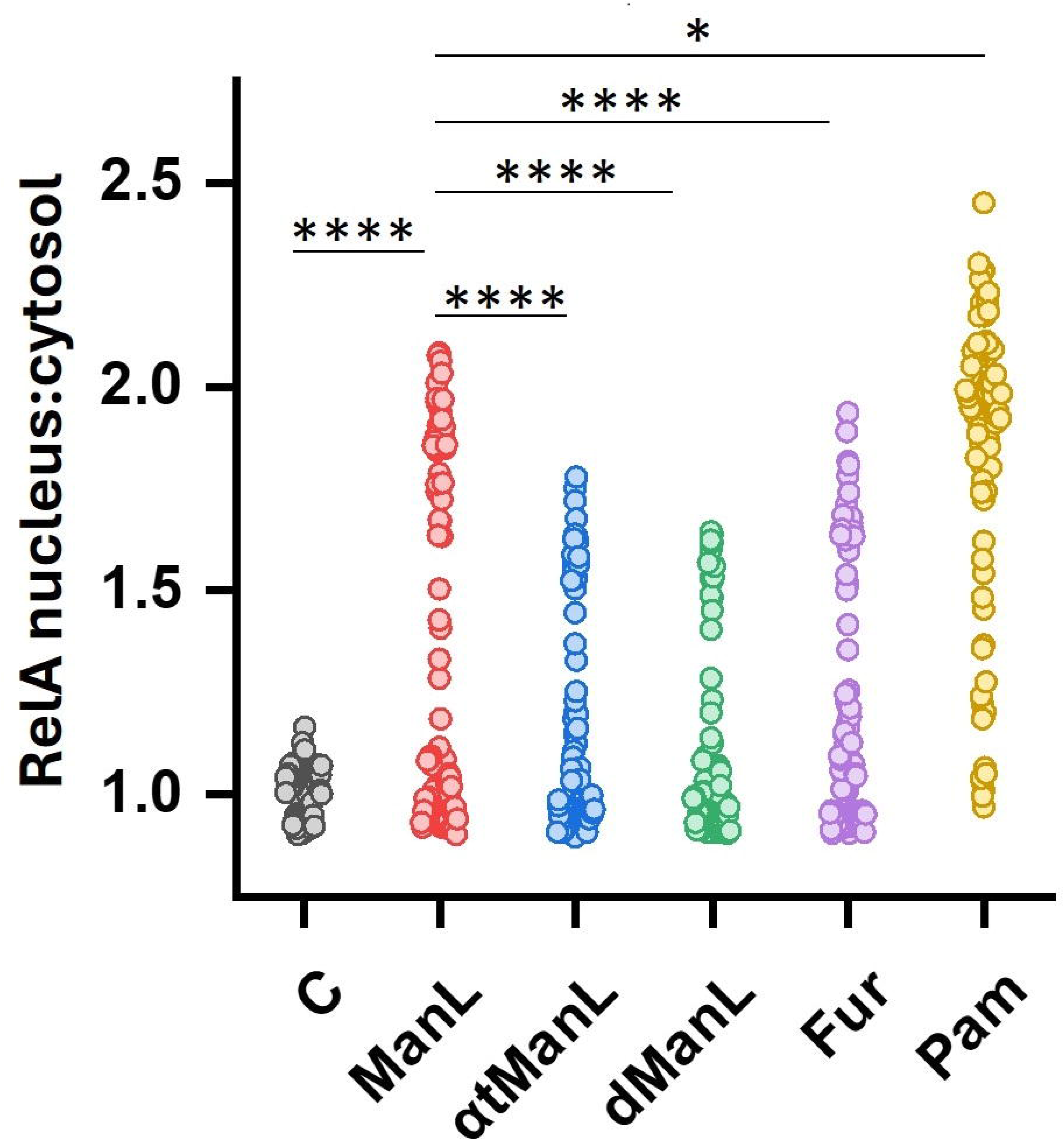

**Figure S5.**
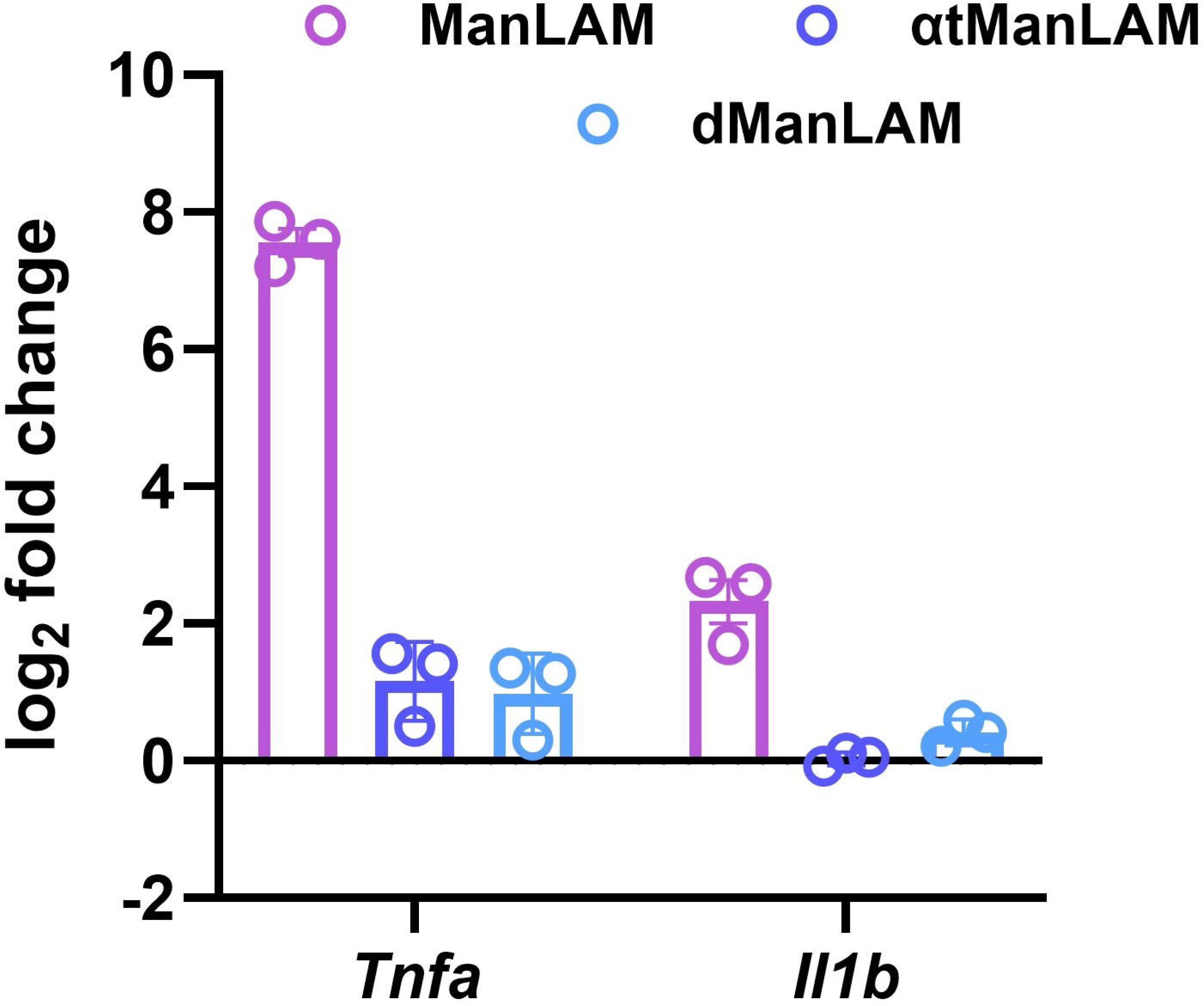

**Figure S6.**
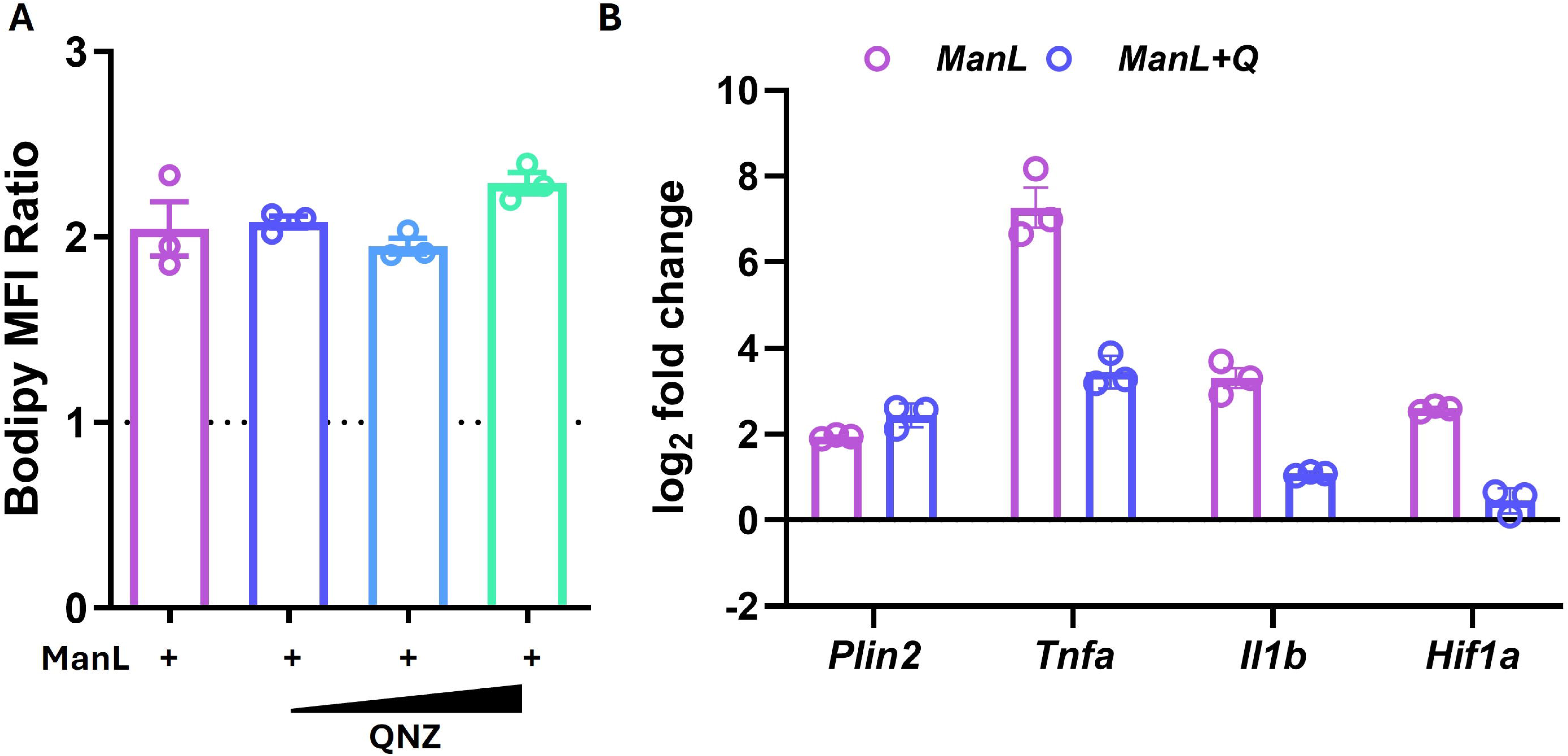

**Figure S7.**
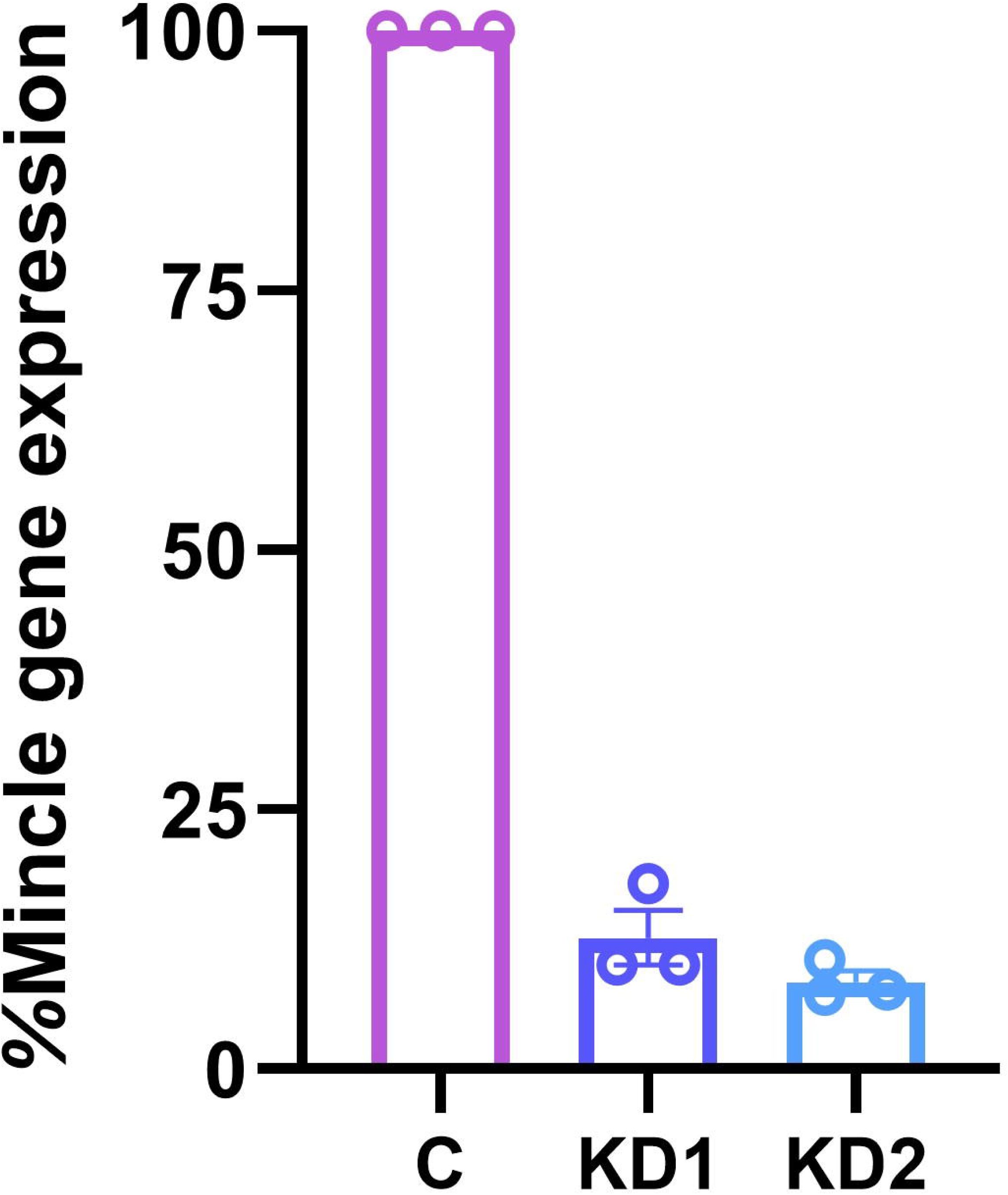

## Notes

### Competing Interest Statement

The authors have declared no competing interest.

### Summary of Updates

Changed Title Changed Abstract Experiments added

