## Supplementary file for "A single mycobacterial ligand organizes multi-receptor signaling to reprogram macrophage lipid metabolism"

**Supplementary Figure Legends**

**Sl Fig. 1: Effects of H_2_O_2_ treatment on *M. tuberculosis* representative lipoprotein LprG.** LprG (Lp; 300 ng/ml, approximately equimolar to ManLAM 500 ng/ml), H_2_O_2_-treated LprG (Lp+H; 300 ng/ml), and vehicle control (C) were tested for ability to induce NF-κB activation in HEK-TLR2 cells. Data were generated and expressed as described in **Fig. 1A**.

**Sl Sl Fig 2.** **Validation of TLR2 and Dectin-2 knock down iBMDM.** **A,D.** Expression levels of TLR2 (A) and Dectin-2 (D) in iBMDM transfected with two different guide RNAs per receptor (KD1and KD2 for TLR2; KD3 and KD4 for Dectin-2) and non-targeting control guide RNAs (C) were measured by qPCR in triplicate and expressed as percent receptor gene expression relative to the housekeeping actin gene. **B,C,E,F.**  Expression levels of downstream genes *Il*6 (**panels B,E**) and *Tnfa* (**panels C,F**) in control (C) KD cells following treatment with the TLR2 agonist Pam3CSK4 (500ng/ml) (panels B,C) or the Dectin-2 agonist furfurman (10 µg/ml) for 2h. Fold induction of *Il*6 and *Tnfa* expression was measured by qPCR in triplicates relative to vehicle-treated cells.

**Sl Fig. 3**. **ManLAM-induced lipid droplet accumulation in iBMDM knocked-down for irrelevant lectin receptors and in human primary macrophages.** Parental control (C), Mincle KD, and Dectin-1 KD iBMDM were treated with 500 ng/ml ManLAM and vehicle for 24 h. Imaging flow cytometry for lipid droplet content were obtained in triplicate and expressed as in **Fig. 1B**. **B.** Human monocyte-derived macrophages were treated for 24h with 500ng/ml ManLAM, demannosylated ManLAM (αtManLAM), and deacylated ManLAM (dManLAM) . Imaging flow cytometry data for lipid droplet content were generated and expressed as in **Fig. 1B**.

**SI Fig. 4.** **Variation of NF-kB activation in RAW 264.7 macrophage cells**. Single-cell scatter plots of NF-κB activation, expressed as nucleus-to-cytosol EGFP-RelA fluorescence intensity ratio, for 200 cells per condition after ligand stimulation for 240 min (ManLAM and derivatives, and furfurman) and 60 min for Pam3CSK4. Each dot represents one cell. Statistical significance was assessed by one-way ANOVA: ***P < 0.001; **P < 0.01; *P < 0.05.

**Sl Fig. 5. Effects on** **ManLAM, αtManLAM and dManLAM on *Tnfa* and *Il1b* gene expression in iBMDM.** Expression levels of *Tnfa* and *Il1b* in iBMDM treated with ManLAM, αtManLAM and dManLAM for 6 hours were measured by RTqPCR in triplicate and expressed as log_2_ fold change of gene expression, relative to the housekeeping actin gene.

**Sl Fig. 6. Effects of the NF-κB inhibitor QNZ on ManLAM-induced lipid droplet accumulation and gene expression. A.** iBMDM were treated with ManLAM (ManL, 500ng/ml) plus vehicle or increasing doses of the NF-κB inhibitor QNZ (5.5, 11, 22nM). Imaging flow cytometry data for lipid droplet content were generated and expressed as in **Fig. 1B**.  B. iBMDM were treated with ManLAM (500ng/ml) plus vehicle (ManL) or QNZ (ManL+Q, 11nM) for 6 hours. Expression levels of *Plin2*, *Tnfa*, *Il1b,* and *Hif1a* were measured by RTqPCR in triplicate and expressed as in the legend to **SI Fig. 5**.

**Sl Fig 7.** **Validation of Mincle knock down iBMDM.** Expression levels of Mincle receptor in iBMDM transfected with two different guide RNAs per receptor (KD1and KD2) and non-targeting control guide RNAs (C) were measured by qPCR in triplicate and expressed as percent receptor gene expression relative to the housekeeping actin gene.

**SI Table 1: qPCR primer sequences**

| Gene name | Primer sequence |
| --- | --- |
| *Tlr2* | F 5’ CACTATCCGGAGGTTGCATATC 3’  R 5’ GGAAGACCTTGCTGTTCTCTAC 3’ |
| *Dectin-2* | F 5’ AATCCCAAGGGAAGGGAGTCTG 3’  R 5’ TGGGCTGGTCCATAATAAATTGG 3’ |
| *Mincle* | F 5’ TGTTTCTTCCAGACAAAGCCCT 3’  R 5’ AATAGAAGTGCTCGTAATGAGTGC 3’ |
| *Tnfa* | F 5’ ATGGCCTCCCTCTCATCAGT 3’  R 5’ GTTTGCTACGACGTGGGCTA 3’ |
| *Il6* | F 5’ ACAAAGCCAGAGTCCTTCAG 3’  R 5’ GTTAGGAGAGCATTGGAAATTGG 3’ |
| *Plin2* | F 5’ GACAGGATGGAGGAAAGACTGC 3’  R 5’ GGTAGTCGTCACCACATCCTTC 3’ |
| *Hif1a* | F 5’ CCTGCACTGAATCAAGAGGTTGC 3’  R 5’ CCATCAGAAGGACTTGCTGGCT 3’ |
| *Il1b* | F 5’ TGTGAAATGCCACCTTTTGA 3’  R 5’ GCTCAAAGGTTTGGAAGCAG 3’ |
| *Actin* | F 5’ GGTGTGATGGTGGGAATGG 3’  R 5’ GCCCTCGTCACCCACATAGGA 3’ |
